# Beyond Core Object Recognition: Recurrent processes account for object recognition under occlusion

**DOI:** 10.1101/302034

**Authors:** Karim Rajaei, Yalda Mohsenzadeh, Reza Ebrahimpour, Seyed-Mahdi Khaligh-Razavi

**Affiliations:** School of Cognitive Sciences (SCS), Institute for Research in Fundamental Sciences (IPM), Niavaran, P.O. Box 19395-5746, Tehran, Iran; Computer Science and AI Lab (CSAIL), MIT, Cambridge, MA, US; Cognitive Science Research Lab., Department of Electrical and Computer Engineering, Shahid Rajaee Teacher Training University, P.O. Box 16785-163, Tehran, Iran; Department of Brain and Cognitive Sciences, Cell Science Research Center, Royan Institute for Stem Cell Biology and Technology, ACECR, Tehran, Iran

## Abstract

Core object recognition, the ability to rapidly recognize objects despite variations in their appearance, is largely solved through the feedforward processing of visual information. Deep neural networks are shown to achieve human-level performance in these tasks, and explain the primate brain representation. On the other hand, object recognition under more challenging conditions (i.e. beyond the core recognition problem) is less characterized. One such example is object recognition under occlusion. It is unclear to what extent feedforward and recurrent processes contribute in object recognition under occlusion. Furthermore, we do not know whether the **conventional** deep neural networks, such as AlexNet, which were shown to be successful in solving core object recognition, can perform similarly well in problems that go beyond the core recognition. Here, we characterize neural dynamics of object recognition under occlusion, using magnetoencephalography (MEG), while participants were presented with images of objects with various levels of occlusion. We provide evidence from multivariate analysis of MEG data, behavioral data, and computational modelling, demonstrating an essential role for recurrent processes in object recognition under occlusion. Furthermore, the computational model with local recurrent connections, used here, suggests a mechanistic explanation of how the human brain might be solving this problem.

**Author Summary:** In recent years, deep-learning-based computer vision algorithms have been able to achieve human-level performance in several object recognition tasks. This has also contributed in our understanding of how our brain may be solving these recognition tasks. However, object recognition under more challenging conditions, such as occlusion, is less characterized. Temporal dynamics of object recognition under occlusion is largely unknown in the human brain. Furthermore, we do not know if the previously successful deep-learning algorithms can similarly achieve human-level performance in these more challenging object recognition tasks. By linking brain data with behavior, and computational modeling, we characterized temporal dynamics of object recognition under occlusion, and proposed a computational mechanism that explains both behavioral and the neural data in humans. This provides a plausible mechanistic explanation for how our brain might be solving object recognition under more challenging conditions.

## 1. Introduction

There is abundance of feedforward, and recurrent connections in the primate visual cortex (Lamme et al., 1998, Sporns and Zwi, 2004). The feedforward connections form a hierarchy of cortical areas along the visual pathway, playing a significant role in various aspects of visual object processing (Felleman and Van Essen, 1991). However, the role of recurrent connections in visual processing have remained poorly understood (Lamme et al., 1998, Lamme and Roelfsema, 2000, Gilbert and Li, 2013, Kafaligonul et al., 2015, Klink et al., 2017).

Several complementary behavioral, neuronal, and computational modeling studies have confirmed that a large class of object recognition tasks called “core recognition” are largely solved through a single sweep of feedforward visual information processing (DiCarlo and Cox, 2007, DiCarlo et al., 2012, Khaligh-Razavi and Kriegeskorte, 2014, Yamins et al., 2014, Cadieu et al., 2014, Wen et al., 2018). Object recognition is defined as the ability to differentiate an object’s identity or category from many other objects having a range of identity-preserving changes (DiCarlo and Cox, 2007). Core recognition refers to the ability of visual system to rapidly recognize objects despite variations in their appearance, e.g. position, scale, and rotation (DiCarlo and Cox, 2007).

Object recognition under challenging conditions, such as high variations (Ghodrati et al., 2014, Karimi-Rouzbahani et al., 2017), object deletion and occlusion (Rensink and Enns, 1998, Nielsen et al., 2006, Wyatte et al., 2012, O’Reilly et al., 2013, Wyatte et al., 2014, Kosai et al., 2014, Choi et al., 2016, Spoerer et al., 2017, Tang et al., 2018), and crowding (Livne and Sagi, 2011, Manassi and Herzog, 2013, Clarke et al., 2014) goes beyond the core recognition problem, which is thought to require more than the feedforward processes. Object recognition under occlusion is one of the key challenging conditions that occurs in many of the natural scenes we interact with every day. How our brain solves object recognition under such challenging condition is still an open question. Object deletion and object occlusion are shown to have different temporal dynamics (Johnson and Olshausen, 2005). While object deletion has been studied before in humans (Nielsen et al., 2006, Wyatte et al., 2012, O’Reilly et al., 2013, Wyatte et al., 2014, Tang et al., 2014, Tang et al., 2018), we do not know much about the dynamics of object processing under the challenging condition of occlusion; in particular there has not been any previous MEG study of occluded object processing, with multivariate pattern analyses approach, linking models with both brain and behavior. Furthermore, as opposed to the core object recognition problem, where the **conventional** feedforward CNNs are shown to explain brain representations (Khaligh-Razavi and Kriegeskorte, 2014, Yamins et al., 2014, Cadieu et al., 2014, Wen et al., 2018), we do not yet have computational models that successfully explain human brain representation and behavior under this challenging condition [Regarding feedforward CNNs, also see (Eberhardt et al., 2016, Rajalingham et al., 2018) where CNNs cannot fully explain patterns of human behavioral decisions].

Few fMRI studies have investigated how and where occluded objects are represented in the human brain (Rauschenberger et al., 2006, Hulme and Zeki, 2007, Hegdé et al., 2008, Ban et al., 2013, Erlikhman and Caplovitz, 2017). Hulme and Zeki (2007) found that faces and houses in fusiform face area (FFA) and lateral occipital cortex (LOC) are represented similary with and without occclusion. Ban et al. (2013) used topographic mapping with simple geometric shapes (e.g. triangles), finding that the occluded portion of the shape is represented topographically in human V1 and V2, suggesting the involvement of early visual areas in object completion. A more recent study showed that the early visual areas may only code spatial information about occluded objects, but not their identity, and higher-order visual areas instead represent object-specific information, such as category or identity of occluded objects (Erlikhman and Caplovitz, 2017). While these studies provide insights about object processing under occlusion, they do not provide any information about the temporal dynamics of these processes, and whether object recognition under occlusion requires recurrent processing.

Our focus in this study is understanding the temporal dynamics of object recognition under occlusion; and whether recurrent connections are critical in processing occluded objects? If yes, in what form are they engaged (e.g. long range feedback or local recurrent?), and how much is their contribution compared to the contribution of the feedforward visual information? We constructed a controlled image set of occluded objects, and used the combination of multivariate pattern analyses (MVPA) of MEG signals, computational modeling, backward masking, and behavioral experiments to characterize representational dynamics of object processing under occlusion, and to determine unique contributions of feedforward and recurrent processes.

Here, we provide five complementary evidence for the contribution of recurrent processes in recognizing occluded objects. *First*, MEG decoding time courses show that onset and peak for occluded objects—without backward masking—are significantly delayed compared to when the whole object is presented without occlusion. The timing of visual information plays an important role in discriminating the processing stages (i.e. feedforward vs. recurrent) with early brain responses reaching higher visual areas being dominantly feedforward and the delayed responses being mainly driven by recurrent processes (Felleman and Van Essen, 1991, Lamme and Roelfsema, 2000, Thorpe and Fabre-Thorpe, 2001, Liu et al., 2009, Khaligh-Razavi et al., 2015, Grootswagers and Carlson, 2015, Kaneshiro et al., 2015, Mohsenzadeh et al., 2018). *Second*, time-time decoding analysis (i.e. temporal generalization) suggests that occluded object processing goes through a relatively long sequence of stages that involve recurrent interaction—likely local recurrent. *Third*, the results of backward masking demonstrate that while the masking significantly impairs both human categorization performances and MEG decoding performances under occlusion, it has no significant effect on object recognition when objects are not occluded. *Fourth*, results from two computational models showed that a **conventional** feedforward CNN (AlexNet) that could achieve human-level performance in the no-occlusion condition, performed significantly worse than humans when objects were occluded. Additionally, the feedforward CNN could only explain the human MEG data when objects were presented without occlusion; but failed to explain the MEG data under the occlusion condition. In contrast, a hierarchical CNN with local recurrent connections (recurrent ResNet) achieved human-level performance and the representational geometry of the model was significantly correlated with that of the MEG neural data when objects were occluded. Finally, we quantified contributions of feedforward and recurrent processes in explaining the neural data, showing a significant unique contribution only for the recurrent processing under occlusion. These findings demonstrate significant involvement of recurrent processes in occluded object recognition, and improve our understand of object recognition beyond the core problem. To our knowledge this is the first MVPA study of MEG data linking feedforward and recurrent deep neural network architectures with both brain and behavior to investigate object recognition under occlusion in humans.

## 2. Results

We used multivariate pattern analysis (MVPA) of MEG data to characterize representational dynamics of object recognition under occlusion (Carlson et al., 2013, Cichy et al., 2014, Isik et al., 2014, Grootswagers et al., 2017). MEG along with MVPA allows for a fine-grained investigation of the underlying object recognition processes across time (Grootswagers et al., 2017, Contini et al., 2017). Subjects (N=15) were presented with images of objects with varying levels of occlusion (i.e., 0% = no-occlusion, 60% and 80% occlusion; Figure 1b). We also took advantage of the visual backward masking (Breitmeyer and Öğmen, 2006) as a tool to further control the feedforward and feedback flow of visual information processing. In the MEG experiment, each stimulus was presented for 34 ms [2 × screen frame rate (17ms) = 34ms], followed by a blank-screen ISI, and then in half of the trials followed by a dynamic mask (Figure S1). We extracted and pre-processed MEG signals from −200 ms to 700 ms with regard to the stimulus onset. To calculate pairwise discriminability between objects, a support vector machine (SVM) classifier was trained and tested at each time point (Figure 1a). MEG decoding time-courses show the pairwise discriminability of object images averaged across individuals. We first present the MEG results of the no-mask trials. After that in section 2.3 we discuss the effect of backward masking.

**Figure 1.**
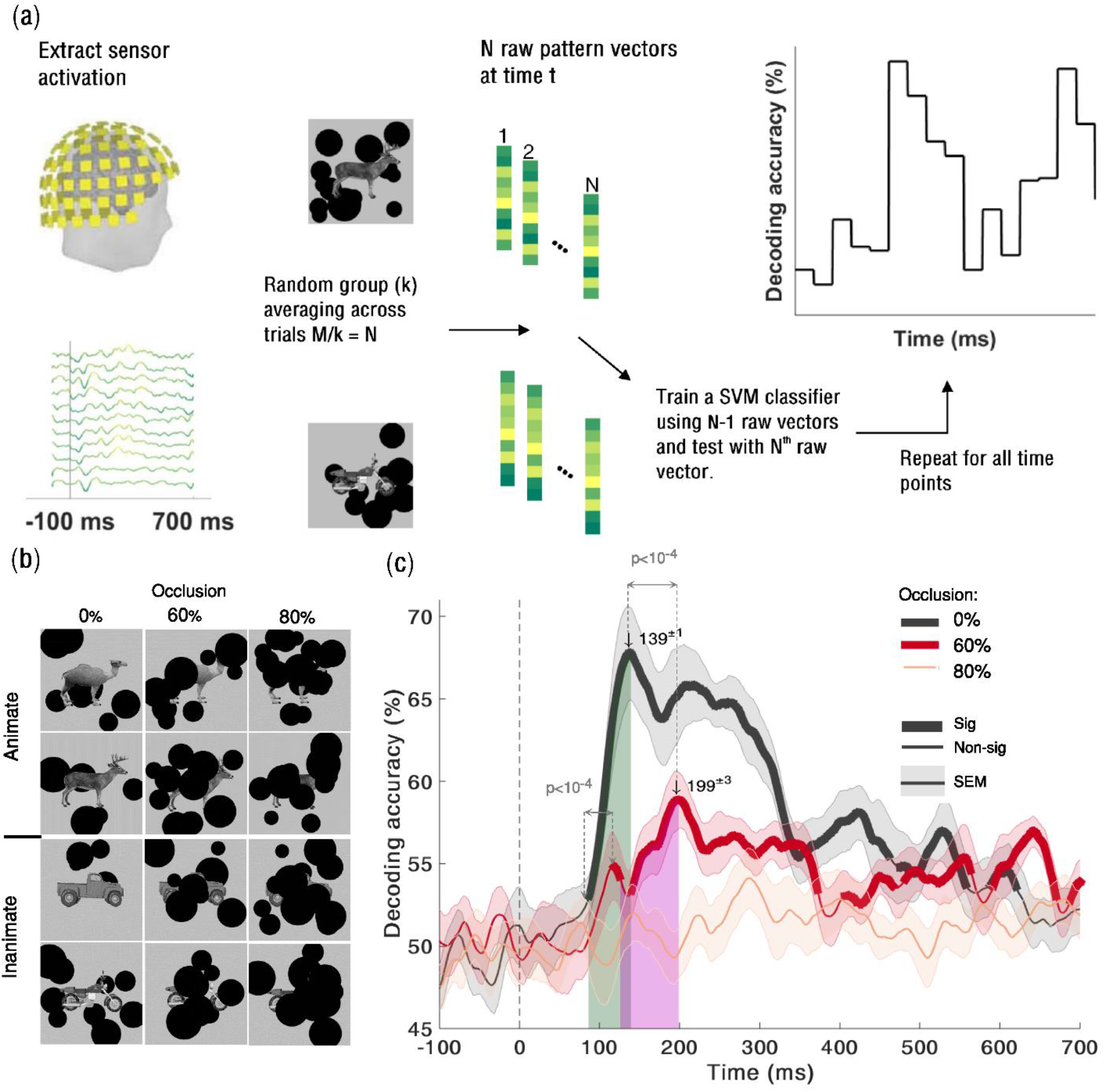
Temporal dynamics object recognition under various levels of occlusion. **(a)** Multivariate pattern classification of MEG data. We extracted MEG signals from −200 ms to 700 ms relative to the stimulus onset. At each time point (ms resolution), we computed average pairwise classification accuracy between all exemplars. **(b**) Sample images of occluded and un-occluded (0% occlusion) objects. There are four object categories: camel, deer, car, and motor. Images are occluded at 0% (no-occlusion), 60%, and 80% occlusion. **(c)** Time courses of pairwise decoding accuracies for the three different occlusion levels (without backward masking) averaged across 15 subjects. Thicker lines indicate a decoding accuracy significantly above chance (right-sided signrank test, FDR corrected across time, p < 0.05), showing that MEG signals can discriminate between object exemplars. Shaded error bars represent standard error of the mean (SEM). The two vertical shaded areas show the time from onset to peak, for 60% occluded and 0% occluded objects, which are largely non-overlapping. The onset latency is 79±3 ms (mean ± SD) in the no-occlusion condition; and 123±15 ms in the 60% occlusion; the difference between onset latencies is significant (p<10^−4^, two-sided signrank test). Arrows above the curves indicate peak latencies. The peak latencies are 139±1ms and 199±3ms for the 0% occluded and partially occluded (60%) objects respectively. The difference between the peak latencies is also statistically significant (p < 10^−4^).

### 2.1. Object recognition is significantly delayed under occlusion

We used pairwise decoding analysis of MEG signals to measure how object information evolves over time (Figure 1a). Significantly above-chance decoding accuracy means that objects can be discriminated using the information available from the brain data at that time-point. The decoding onset latency indicates the earliest time that the object-specific information becomes available and the peak decoding latency is the time-point wherein we have the highest object-discrimination performance.

We found that object information emerges significantly later under occlusion compared to the no-occlusion condition. Object decoding under no-occlusion had an early onset latency at 79ms [±3 ms standard deviation (SD)] and was followed by a sharp increase reaching its maximum accuracy (i.e. peak latency) at 139±1 ms (Figure 1c). This early and rapidly evolving dynamic is well consistent with the known time-course of the feedforward visual object processing [see Figure S2 and (Liu et al., 2009, Carlson et al., 2013, Cichy et al., 2014)].

However, when objects were partially occluded (i.e. 60% occlusion), decoding time-courses were significantly slower than the 0% occlusion condition: the onset for decoding accuracy was at 123±15 ms followed by a gradual increase in decoding accuracy until it reached its peak decoding accuracy at 199±3 ms (Figure 1c). The difference between onset latencies and peak latencies were both statistically significant with p<10^−4^ (two sided sign-rank test). Analysis of the behavioral response times was also consistent with the MEG object decoding curves, showing a significant delay in participants’ response times when increasing the occlusion level (Figure S3). The slow temporal dynamics of object recognition under occlusion and the observed significant temporal delay in processing occluded objects compared to un-occluded (0% occlusion) objects do not match with a fully feedforward account of visual information processing. Previous studies have shown that first responses to visual stimuli that contain category-related information can reach higher visual areas in as little as 100 ms. (Thorpe, 2009, Liu et al., 2009, Cichy et al., 2016b). Therefore, the observed late onset and the significant delay in peak and onset of the decoding accuracy for occluded objects, may be best explained by the engagement of recurrent processes.

Under 80% occlusion, the MEG decoding results did not reach significance [right-sided signrank test, FDR corrected across time, p < 0.05] (Figure 1c). However, behaviorally, human subjects still performed above-chance in object categorization even under 80% occlusion (Figure 5d). This discrepancy might be due to MEG acquisition noise, whereas the behavioral categorization task is by definition free from that type of noise.

While the MEG and behavioral data have different levels of noise, we showed that within the MEG data itself, presented images with different levels of occlusion (0%, 60%, 80%) did not differ in terms of their level of MEG noise (Figure S4). Thus, the difference in decoding performance between different levels of occlusion cannot be simply explained by the difference in noise. Furthermore, patterns of cross-decoding analyses (Figure 3) demonstrates that the observed delay in peak latency under occlusion cannot be simply explained by a difference in signal strength.

### 2.2. Time-time decoding analysis for occluded objects suggests a neural architecture with recurrent interactions

We performed time-time decoding analysis measuring how information about object discrimination generalizes across time (Figure 2a). Time-time decoding matrices are constructed by training a SVM classifier at a given time point and testing its generalization performance at all other time-points (see Methods). The pattern of temporal generalization provides useful information about the underlying processing architecture (King and Dehaene, 2014).

**Figure 2.**
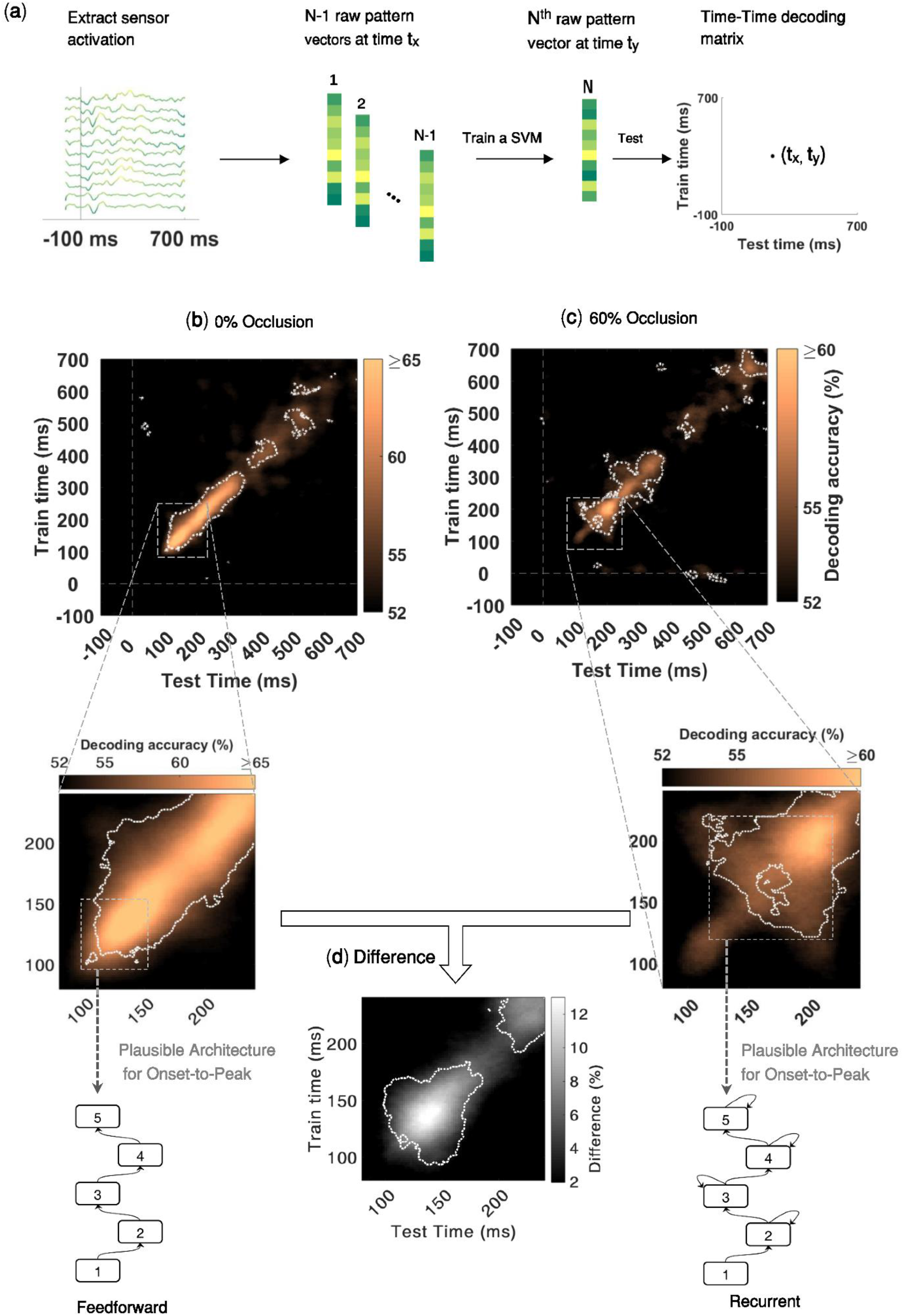
Temporal generalization patterns of object recognition with and without occlusion. **(a)** Time-time decoding analysis. The procedure is similar to the calculations of pairwise decoding accuracies explained in Figure 1, except that here the classifier is trained at a given time-point, and then tested at all time-points. In other words, for each pair of time points (t_x_, t_y_), a SVM classifier is trained by N-1 MEG pattern vectors at time t_x_ and tested by the remaining one pattern vector at time t_y_, resulting to an 801×801 time-time decoding matrix. (**b-c**) Time-time decoding accuracy and plausible processing architecture for no-occlusion and 60% occlusion. The results are for MEG trials without backward masking. Horizontal axis indicates testing times and vertical axis indicates training times. Color bars represent percent of decoding accuracies (chancel level = 50%); please note that in the time-time decoding matrices, the color bar ranges for 0% occlusion and 60% occlusion are different. Within the time-time decoding matrices, significantly above chance decoding accuracies, are surrounded by the white dashed contour lines (right-sided signrank test, FDR corrected across the whole 801×801 decoding matrix, p < 0.05). For each time-time decoding matrix, we also show the plausible processing architecture corresponding to it. These are derived from the observed patterns of temporal generalization from onset-to-peak decoding (shown by the gray dashed rectangles) [see Figure 5 of (King et al., 2016)]. Generalization patterns for the no-occlusion condition are consistent with a hierarchical feedforward architecture; whereas, for the occluded objects (60%) the temporal generalization patterns are consistent with a hierarchical architecture with recurrent connections. **(d)** Difference in time-time decoding accuracies between no-occlusion and occlusion conditions. Significantly above zero differences are surrounded by the white dashed contour lines (right-sided signrank test, FDR corrected across [80-240]ms matrix at p < 0.05).

We were interested to see if there are differences between temporal generalization patterns of occluded and un-occluded objects. Different processing dynamics may lead to distinct patterns of generalization in the time-time decoding matrix [see (King and Dehaene, 2014) for a review]. For example, a narrow diagonal pattern suggests a hierarchical sequence of processing stages wherein information is sequentially transferred between neural stages. This hierarchical architecture is well consistent with the feedforward account of neural information processing across the ventral visual pathway. On the other hand, a time-time decoding pattern with off-diagonal generalization suggests a neural architecture with recurrent interactions between processing stages [see Figure 5 in (King et al., 2016)].

The temporal generalization pattern under no-occlusion (Figure 2b) indicated a sequential architecture, without an off-diagonal generalization until its early peak latency at 140 ms. This is consistent with a dominantly feedforward account of visual information processing. There was some off-diagonal generalization after 140 ms, however that was not of interest here, because the ongoing recurrent activity after the peak latency (as shown in Figure 1c) did not carry any information that further improves pairwise decoding performance of 0% occluded objects. On the other hand, when objects were occluded, the temporal generalization matrix (Figure 2c) indicated a significantly delayed peak latency at 199ms with extensive off-diagonal generalization before reaching its peak. In other words, for occluded objects, we see a discernible pattern of temporal generalization, which is characterized by 1) a relatively weak early diagonal pattern of the decoding accuracy during [100 150]ms with limited temporal generalization, which is in contrast with the high accuracy decoding of 0% occluded objects in the same time period. 2) A relatively late peak decoding accuracy with a wide generalization pattern around 200ms. This pattern of temporal generalization can be simulated by a hierarchical neural architecture with local recurrent interactions within the network [Figure 5 of (King et al., 2016)]

We also performed sensorwise decoding analysis to explore spatio-temporal dynamics of object information. To calculate sensorwise decoding, pairwise decoding analysis was conducted on 102 neighboring triplets of MEG sensors (2 gradiometers and 1 magnetometer in each location) yielding a decoding map of brain activity at each time-point. The sensorwise decoding patterns indicated the approximate locus of neural activity: in particular, we see that for both 0% occlusion (supp. movie 1) and 60% occlusion (supp. movie2) conditions, during the onset of decoding as well as the peak decoding time, the main source of object decoding is in the left posterior-temporal sensors. From [110ms to 200ms], the peak of decoding accuracy remains locally around the same sensors, suggesting a sustained local recurrent activity.

**Generalization across time and occlusion levels:** Time-time decoding analyses can be further expanded by training the classifiers in one condition (e.g. occluded) and testing their ability to generalize to the other condition (e.g. un-occluded). The resulting pattern of generalization across time and occlusion level provides diagnostic information about the organization of brain processes (Figure 3). In particular, this can provide us with further evidence as to whether the observed decoding results under occlusion is due to changes in activation intensity (e.g. weakening of the signal), or a genuine difference in latency. As shown in ((King and Dehaene, 2014), Figure 4) each of these two come with distinct cross-condition generalization matrices. A change of signal intensity leads to asymmetric generalizations across conditions because it can be more efficient to train a decoder with relatively high signal-to-noise ratio (SNR) data (e.g. 0% occlusion) and generalize it to low SNR data (e.g. 60% occlusion) rather than vice versa. However, in Figure 3, we do not see such asymmetry in decoding strengths, instead we observe different generalization patterns that are more consistent with changes in the speed of information processing (i.e. latency). More specifically, when the classifier is trained with 0% occlusion and tested on 60% occlusion (upper-left matrix in Figure 3a) the generalization pattern is shifted to the above diagonal and vice versa when trained with 60% occlusion and tested on 0% occlusion (the lower-right matrix).

**Figure 3.**
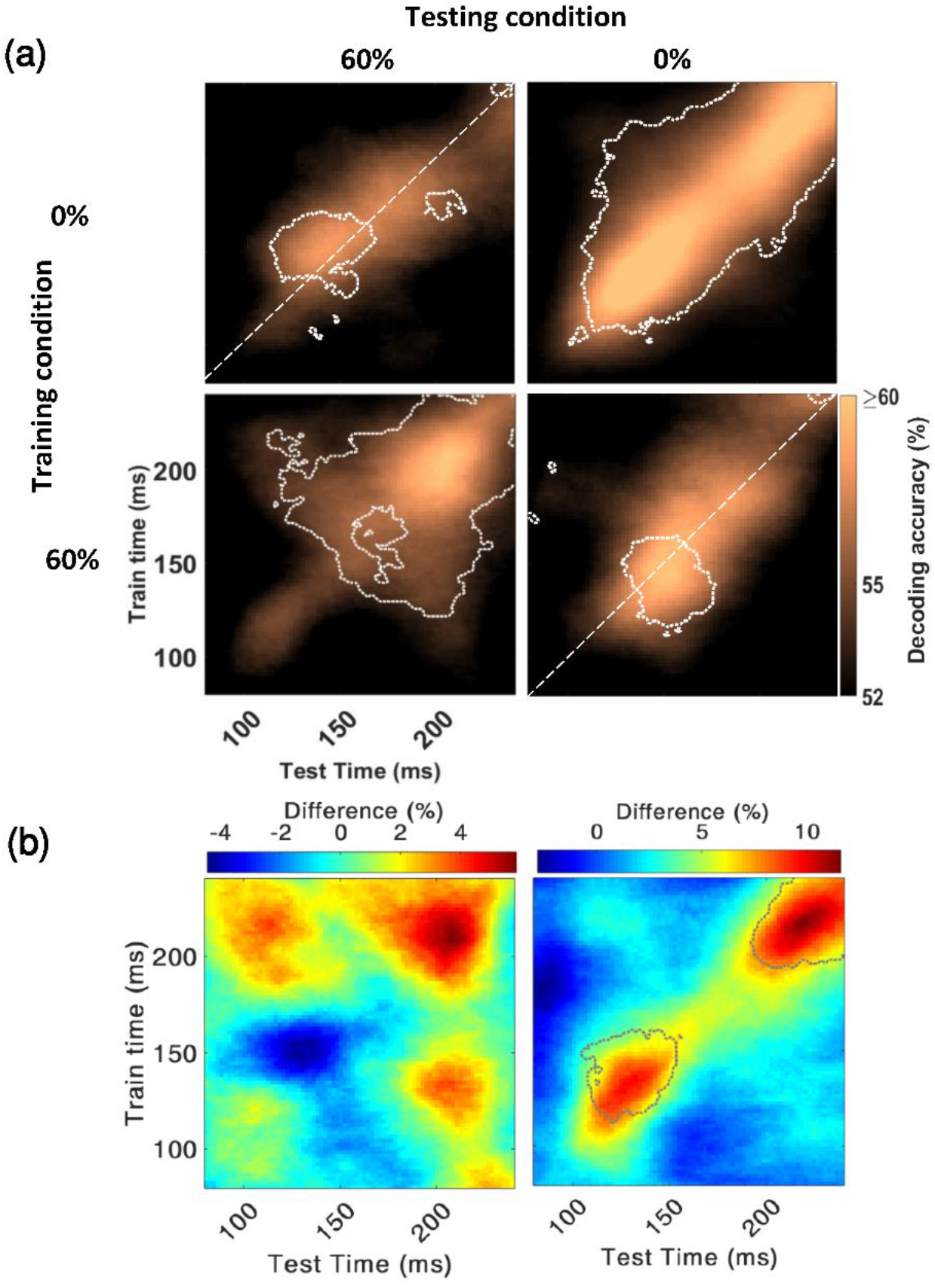
Generalization across time and occlusion levels. **(a)** The classifier is trained on an occlusion level (e.g. 0% occlusion) and tested on the other occlusion level (e.g. 60% occlusion). Time-points with significant decoding accuracy are shown inside the dashed contours (right-sided signrank test, FDR-corrected across time, p<0.05). The contour of significant time-points has a shift towards the upper side of the diagonal when the classifier is trained with 0% occlusion and tested on 60% occlusion (i.e. 63% of significant time points are above the diagonal) whereas in the lower right matrix we see the opposite pattern (66% of significant time points are located below the diagonal). **(b)** The two color maps below the decoding matrices show the difference between the two decoding matrices located above them.

**Figure 4.**
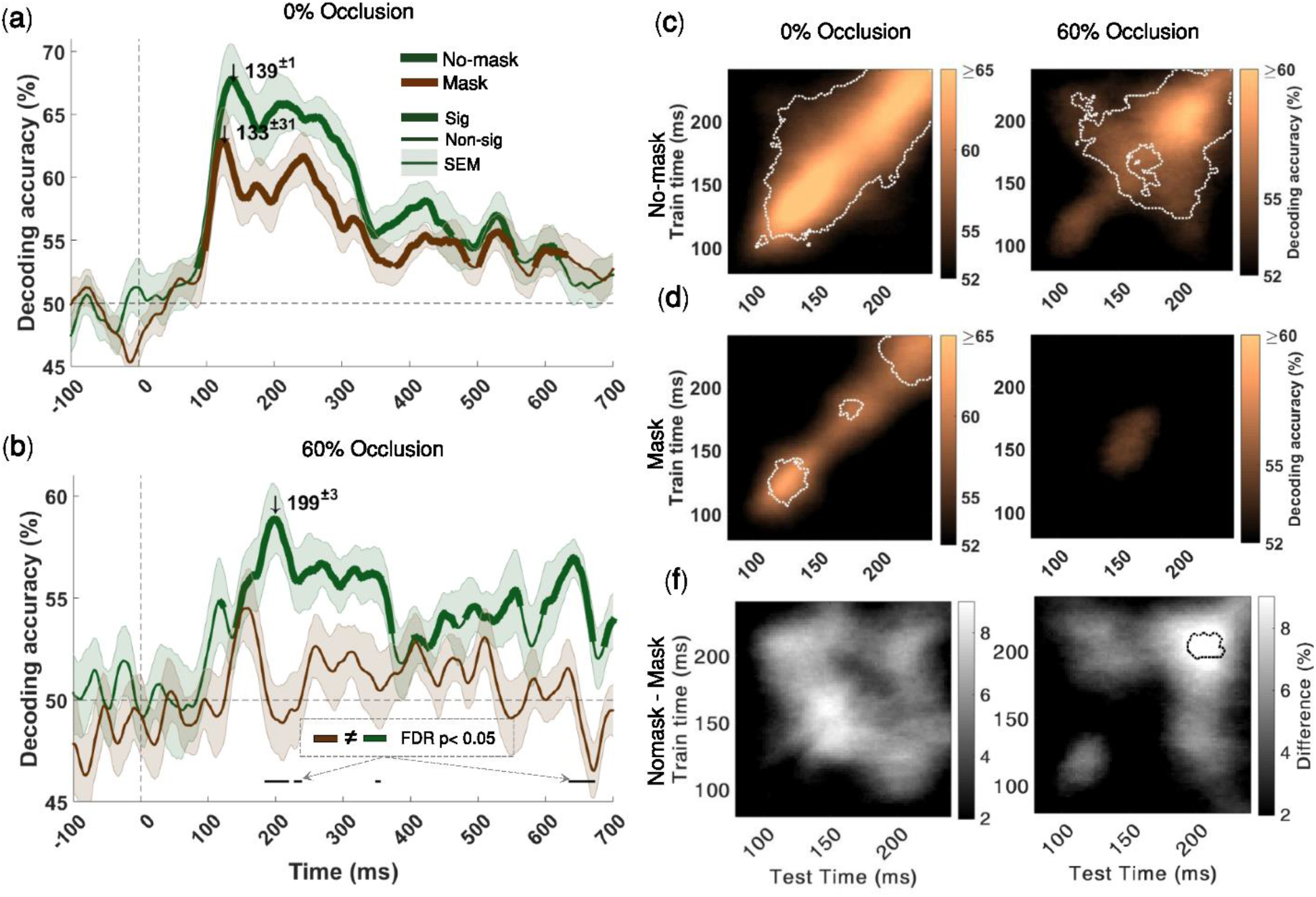
Backward masking significantly impairs object decoding under occlusion, but has no significant effect on object decoding under no occlusion. (a) Time-courses of the average pairwise decoding accuracies under no-occlusion. Thicker lines indicate significant time-points (right-sided signrank test, FDR corrected across time, p < 0.05). Shaded error bars indicate SEM (standard error of the mean). Downward pointing arrows indicate peak decoding accuracies. There is no significant difference between decoding time-courses for mask and no-mask trials, under no-occlusion **(b)** Time-courses of the average pairwise decoding under 60% occlusion (for 80% occlusion see Figure S5). Under occlusion, the decoding onset latency for the no-mask trials is 123±15ms, with its peak decoding accuracy at 199±3ms; whereas the time-course for the masked trials does not reach statistical significance, demonstrating that backward masking significantly impairs object recognition under occlusion. Black horizontal lines below the curves show the time-points at which the two decoding curves are significantly different. This is particularly evident around the peak latency of the no-mask trials [from 185ms to 237ms]. **(c, d)** Time-time decoding matrices of 60% occluded and (0%) un-occluded objects with and without backward masking. Horizontal axes indicate testing times and the vertical axes indicate training times. Color bars show percent of decoding accuracies. Please note that in the time-time decoding matrices, the color bar ranges for 0% occlusion and 60% occlusion are different. Significantly above chance decoding accuracies, are surrounded by the white dashed contour lines (right-sided signrank test, FDR corrected across the whole 801×801 decoding matrix, p < 0.05). **(f)** Difference between time-time decoding matrices with and without backward masking. Statistically significant differences are surrounded by the black dotted contours (right-sided signrank test, FDR corrected across time at p < 0.05). There are significant differences between mask and no-mask only under occlusion.

### 2.3. Backward masking significantly impaired object recognition only under occlusion

Visual backward masking has been used as a tool to disrupt the flow of recurrent information processing, while feedforward processes are left relatively intact (Lamme and Roelfsema, 2000, Lamme et al., 2002, Bacon-Macé et al., 2005, Breitmeyer and Öğmen, 2006, Fahrenfort et al., 2007, Serre et al., 2007, Ghodrati et al., 2014). Our time-time decoding results (Figure 4d 0% occluded) additionally supports the recurrent explanation of backward masking: off-diagonal generalization in time-time decoding matrices are representative of recurrent interactions; these off-diagonal components disappear when backward masking is present.

Considering the recurrent explanation of the masking effect, we further examined how the recurrent processes contribute in object processing under occlusion. We found that backward masking significantly reduced both MEG decoding accuracy time-course (Figure 4b) and subjects’ behavioral performances (Figure 5d), only when objects were occluded. When occluded objects are masked, the MEG decoding time-course from 185ms to 237ms is significantly lower than the decoding time-course when in no-mask condition (Figure 4b, black horizontal lines; two-sided signrank test, FDR-corrected across time p < 0.05). On the other hand, for un-occluded objects, we did not find any significant difference between decoding time-courses of the mask and no-mask conditions (Figure 4a).

**Figure 5.**
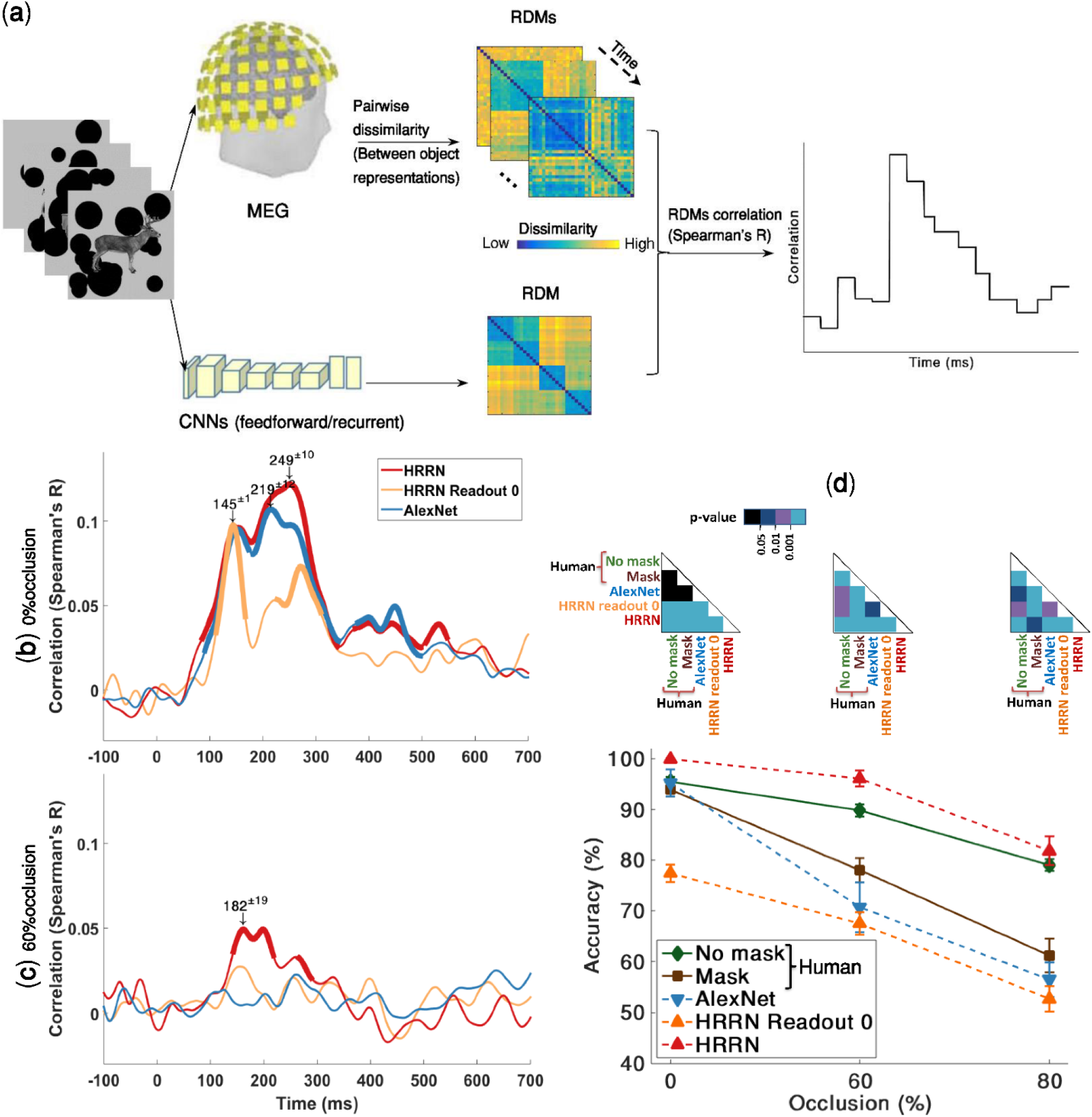
Comparing human MEG and behavioral data with feedforward and recurrent computational models of visual hierarchy. **(a)** Time-varying representational similarity analysis between human MEG data and the computational models. We, first, obtained representational dissimilarity matrices (RDM) for each computational model —using feature values of the layer before the softmax operation—, and for the MEG data at each time-point. For each subject, their MEG RDMs were correlated (Spearman’ R) with the computational model RDMs (i.e. AlexNet & HRRN) across time; the results were then averaged across subjects. **(b, c)** Time-courses of RDM correlations between the models and the human MEG data. HRRN readout stage 0 represents the purely feedforward version of HRRN. Thicker lines show significant time points (right-sided signrank test, FDR-corrected across time, p <= 0.05). We indicate peak correlation latencies by numbers (mean ± SD) above the downward pointing arrows. Under no-occlusion, AlexNet and HRRN demonstrate almost similar time-courses except that the peak latency for HRRN (249±10ms) is significantly later than the peak latency for AlexNet (219±12ms). However, under occlusion, only HRRN showed significant correlation with MEG data, with a peak latency of 182±19ms. **(d)** Object recognition performance of humans (mask and no-mask trials) and models [AlexNet and HRRN-ReadoutStage-0 (feedforward) and HRRN(recurrent)] across different levels of occlusion. We evaluated model accuracies on a multiclass recognition task similar to the multiclass behavioral experiment done in humans (Figure S6). The models’ performances were calculated by holding out an occlusion level for testing, and training a SVM classifier on the remaining levels of occlusion. Error bars are SEM.

Consistent with the MEG decoding results, while the masking significantly reduced behavioral categorization performances when objects were occluded, it had no significant effect on the categorization performance for the un-occluded objects (Figure 5d) [two-sided signrank test]. Particularly, the backward masking removed the late MEG decoding peak (around 200ms) under occlusion (Figure 4f) likely due to disruption of later recurrent interactions.

Taken together, we demonstrated that visual backward masking, which is suggested to disrupt recurrent processes (Lamme and Roelfsema, 2000, Lamme et al., 2002, Breitmeyer and Öğmen, 2006, Fahrenfort et al., 2007, Macknik and Martinez-Conde, 2007), significantly impairs object recognition only under occlusion. On the other hand, masking did not affect object processing under no occlusion, when information from the feedforward sweep is shown to be sufficient for object recognition. Thus, providing further evidence for the essential role of recurrent processes in object recognition under occlusion.

### 2.4. How well does a computational model with local recurrent interactions explain neural and behavioral data under occlusion?

Recent studies have shown that convolutional neural networks (CNNs) achieve human-level performance and explain neural data under non-challenging conditions—also referred to as the core object recognition (DiCarlo and Cox, 2007, Khaligh-Razavi and Kriegeskorte, 2014, Yamins et al., 2014). Here, we first examined whether such feedforward CNNs (i.e. AlexNet) can explain the observed human neuronal and behavioral data in a challenging object recognition task when objects are occluded. The model accuracy was evaluated by the same object recognition task used to measure human behavioral performance (Figure S6). A multiclass linear SVM classifier was trained on images from two occlusion levels and tested on the left-out occlusion level, using features from the penultimate layer of the model (e.g. ‘fc7’ in AlexNet). The classification task was to categorize images into car, motor, deer, or camel. This procedure was repeated for 15 times, and the mean categorization performance is reported here.

We, additionally, used representational similarity analysis (RSA) to assess model’s performance in explaining the human MEG data. RSA correlates time-resolved human MEG representations with that of the model, on the same set of stimuli. First, dissimilarity matrices (RDMs) were separately calculated for the MEG signals and the model. The model RDM were then correlated with MEG RDMs across time (Figure 5a; also see Methods).

We found that in the no-occlusion condition, the feedforward CNN achieved human-level performance and CNN representations were significantly correlated with the MEG data. Significant correlation between the model and MEG representational dissimilarity matrices (RDMs) started at ∼90ms after the stimulus onset and remained significant for several hundred milliseconds with two peaks at 150ms and 220ms (Figure 5b). However, the feedforward CNNs (i.e. AlexNet and the purely feedforward version of HRRN (HRRN with readout stage 0)) failed to explain human MEG data when objects were occluded. And the model performance was significantly lower than that of human in the occluded object recognition task.

We were wondering if a model with local recurrent connections could account for object recognition under occlusion. Inspired by recent advancements in deep convolutional neural networks (He et al., 2016a, He et al., 2016b, Liao and Poggio, 2016, Veit et al., 2016), we built a hierarchical recurrent ResNet (HRRN) that follows the hierarchy of the ventral visual pathway (Figure 6, also see Methods for more details about the model). The recurrent model (HRRN) could rival the human performance in the occluded object recognition task (Figure 5d), performing strikingly better than AlexNet in 60% and 80% occlusion. We also compared confusion matrices (patterns of errors) between the models and human (Figure S7). Under the no-mask condition, HRNN had a significantly higher correlation with humans under 0% and 80% occlusion (the difference was not significant in 60% occlusion, Figure S7b).

**Figure 6.**
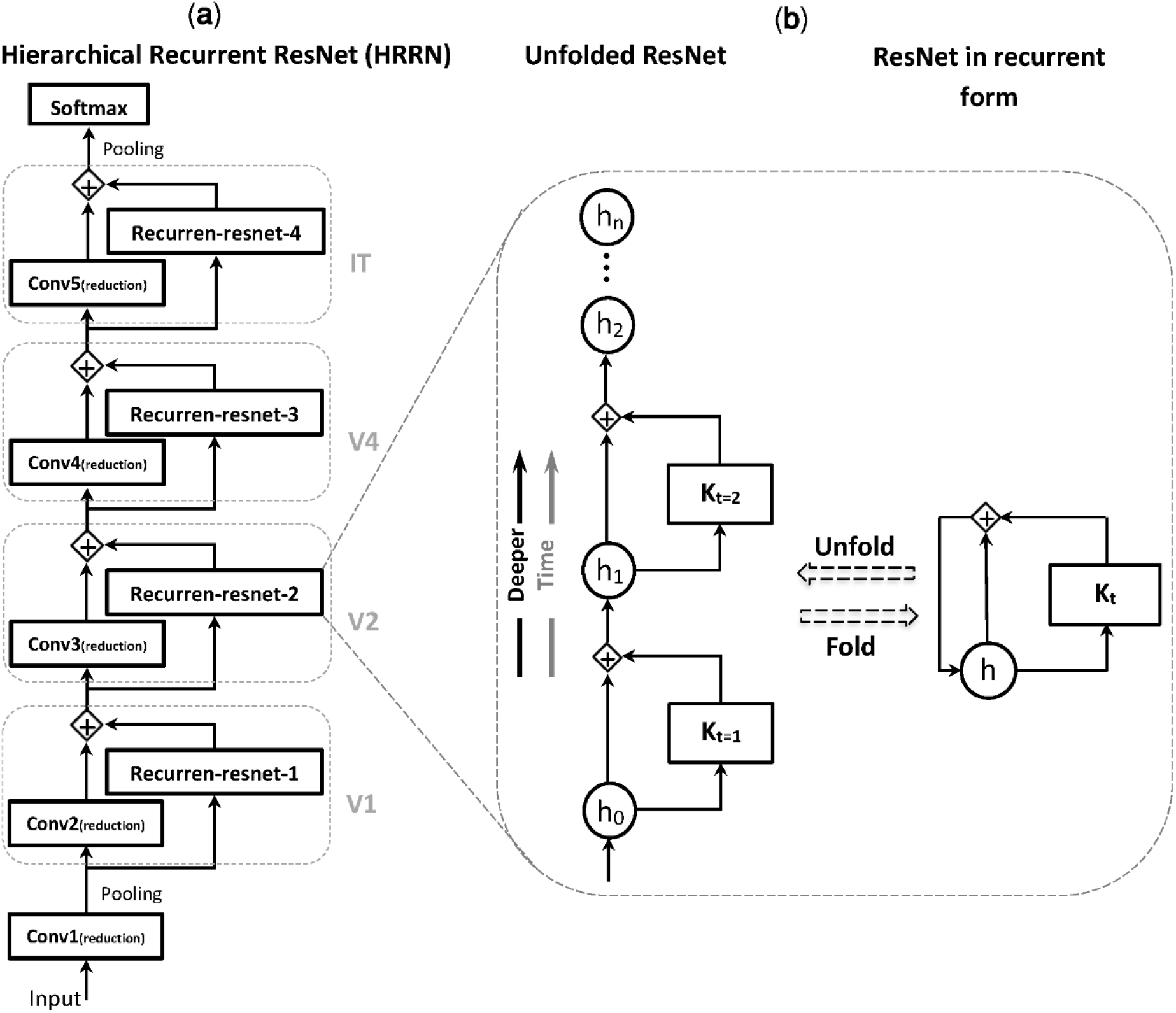
Hierarchical Recurrent ResNet (HRRN) in unfolded form is equivalent to an ultra-deep ResNet. **(a)** A hierarchy of convolutional layers with local recurrent connections. This hierarchical structure models the feedforward and local recurrent connections along the hierarchy of ventral visual pathway (e.g. V1, V2, V4, IT). **(b)** Each recurrent unit is equivalent to a deep ResNet with arbitrary number of layers depending on the unfolding depth. *h*_t_ is the layer activity at a specific time (t) and K_t_ represents a sequence of nonlinear operations (e.g. convolution, batch normalization, and ReLU). [see (Liang and Hu, 2015) for more info]

Additionally, the HRRN model representation was significantly correlated with that of the human MEG data under occlusion [Figure 5c] (onset = 138±2ms; peak = 182±19ms). It is worth noting that the recurrent model here may be considered functionally equivalent to an arbitrarily deep feedforward CNN. We think the key difference between AlexNet and HRRN, is indeed in the number of non-linear transformations applied to the input image (please refer to section 3.3 for a discussion about this).

The models here, the purely feedforward models (i.e. HRRN readout stage 0, and AlexNet) and the model with local recurrent connections, were both trained on the same object image dataset [ImageNet (Deng et al., 2009)] and had equal number of feedforward convolutional layers. Both models performed similarly in object recognition under no-occlusion, and achieved human-level performance. However, under occlusion, only the HRRN (i.e. the model with recurrent connections) could partially explain the human MEG data and achieved human-level performance, whereas the purely feedforward models failed to achieve human-level performance under occlusion—in both MEG and behavior.

### 2.5 Contribution of feedforward and recurrent processes in solving object recognition under occlusion

To quantify the contribution of feedforward and recurrent processes in solving object recognition under occlusion, we first correlated the feedforward and recurrent model RDMs with the average MEG RDMs extracted from two time spans: 80 to 150 ms, which is dominantly feedforward (Liu et al., 2009, Cichy et al., 2016b, Mohsenzadeh et al., 2018), and 151 to 300 ms (significant involvement of recurrent processes). The results are shown in Figure 7a. HRRN and AlexNet both have a similar correlation with the MEG data at [80-150 ms]. However, the HRRN shows a significantly higher correlation with the MEG data at [151-300 ms] compared to the AlexNet.

**Figure 7.**
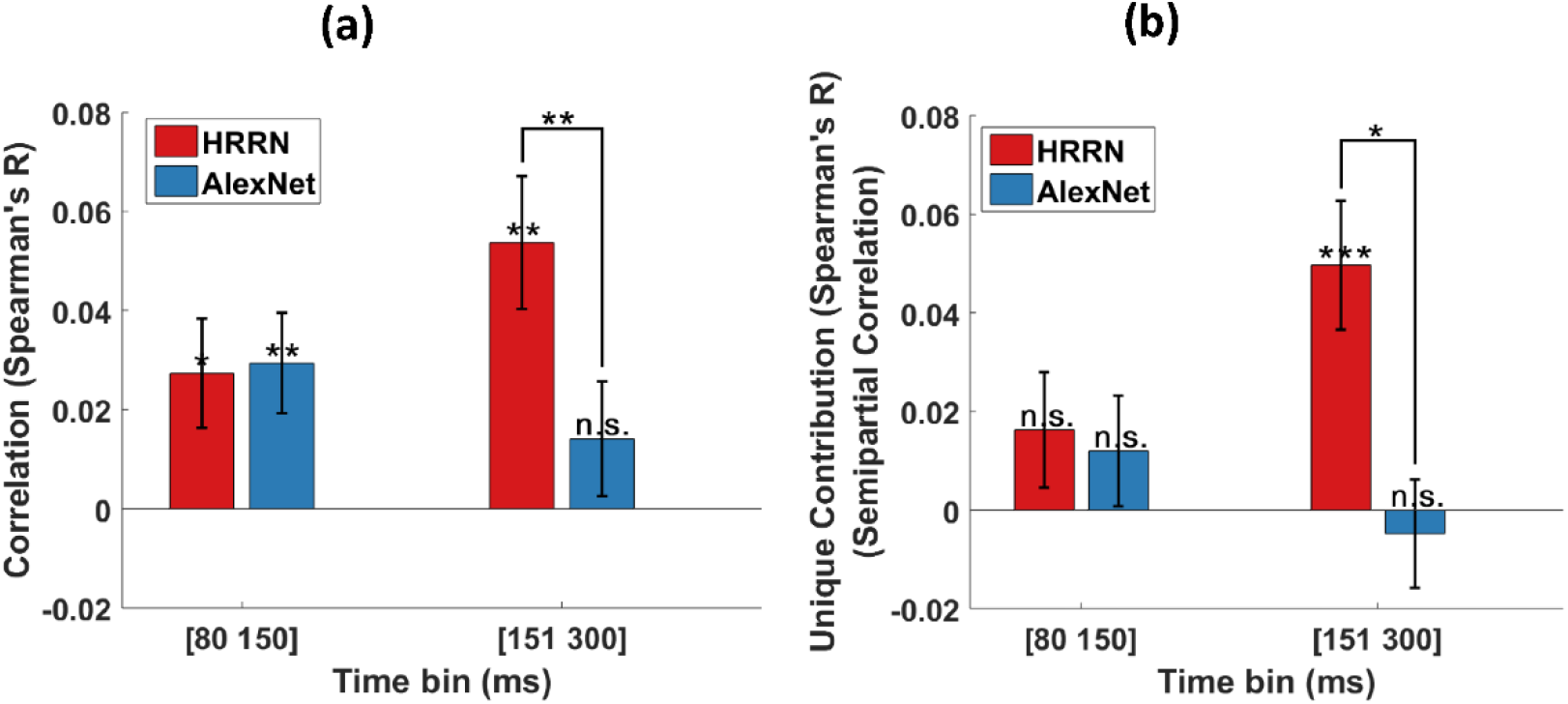
Contribution of the feedforward and recurrent models in explaining MEG data under 60% occlusion. **(a)** Correlation between the models RDMs and the average MEG RDM over two different time bins. **(b)** Unique contribution of each model (semipartial correlation) in explaining the MEG data. Error bars represent SEM (Standard Error of the Mean). Significantly above zero correlations/semipartial-correlations and significant differences between the two models are indicated by stars. * = p<0.05; ** = p<0.01; *** = p<0.001.

We were further interested to determine the unique contribution of the models in explaining the neural data under occlusion. To this end, we calculated semipartial correlations between the model RDMs and the MEG RDMs (Figure 7b). We find that the HRRN and AlexNet perform similarly in explaining the mainly feedforward component (i.e. 80-150ms) of the MEG data and they do not have a significant unique contribution. On the other hand, for the later component (150-300 ms) of the MEG data, only the HRRN model has a unique contribution.

## 3. Discussion

We investigated how the human brain processes visual objects under occlusion. Using multivariate MEG analysis, behavioral data, backward masking and computational modeling, we demonstrated that recurrent processing plays a major role in object recognition under occlusion.

One of the unique advantages of the current work in comparison to a number of previous studies that have investigated object recognition under partial visibility (Wyatte et al., 2014, Tang et al., 2014) was bringing together neural data, behavioral data and computational models in a single study. This enabled us to study the link between brain and behavior and propose a neural architecture that is consistent with both. Whereas previous studies either related models with behavior (e.g. (Wyatte et al., 2014)), missing the link with brain data; or otherwise highlighting the role of recurrent processes using neural data (Tang et al., 2014) without suggesting a computational mechanism that could explain the data and the potential underlying mechanisms (however, please also see the recently published (Tang et al., 2018)).

Another unique advantage of the current study was the comparison of different deep neural network architectures and comparing their ability in explaining neural and behavioral data under occlusion. To our knowledge this was the first work that specifically compared deep convolutional networks with neural data in occluded object processing.

### 3.1. Beyond core object recognition

Several recent findings have indicated that a large class of object recognition tasks referred to as ‘core object recognition’ are mainly solved in the human brain within the first ∼100 ms after stimulus onset (Thorpe, 2009, Liu et al., 2009, Carlson et al., 2013, Isik et al., 2014, Cichy et al., 2016a), largely associated with the feedforward path of visual information processing (Lamme and Roelfsema, 2000, Khaligh-Razavi and Kriegeskorte, 2014, Yamins et al., 2014, Cadieu et al., 2014). More challenging tasks, such as object recognition under occlusion, go beyond the core recognition problem. So far it has been unclear whether the visual information from the feedforward sweep can fully account for this or otherwise recurrent information are essential to solve object recognition under occlusion.

#### 3.1.1. Temporal dynamics

We found that under the no-occlusion condition, the MEG object-decoding time-course peaked at 140ms with an early onset at 79ms, consistent with findings from previous studies (DiCarlo and Cox, 2007, Carlson et al., 2013, Cichy et al., 2014, Isik et al., 2014). Intracranial recordings in human visual cortex have shown that early responses to visual stimuli can reach higher visual areas in as little as 100 ms, with a peak decoding performance after 150 ms (Liu et al., 2009, Cichy et al., 2016b). These results suggest that category-related information around ∼100 ms (±20 ms) after stimulus onset are mainly driven by feedforward processes (see Figure S2 for time course of visual processing in human). M/EEG studies in humans have additionally shown that approximately after 150 ms, a very complex and dynamic phase of visual processing may strengthen the category-specific semantic representations (Clarke et al., 2011, Clarke, 2015), which are likely to be driven by recurrent (Thorpe, 2009). In our study, when objects were occluded, object decoding accuracy peaked at 200ms, with a late onset at 123ms—significantly later than the peak and onset under the no-occlusion condition (i.e. 140ms and 79ms)—suggesting the involvement of recurrent processes. Given the results from the temporal generalization analysis (Figure 2c), and the computational modelling, we argue for the engagement of mostly *local* recurrent connections as opposed to long-range top-down feedback in solving object recognition under occlusion for this image set. Previous studies also suggest that long-range top-down recurrent (e.g. PFC to IT) is prominently engaged after 200ms from stimulus onset (Tomita et al., 1999, Garrido et al., 2007, Liu et al., 2009, Goddard et al., 2016). However, we do not rule out the possibility of involvement of long-range feedback in processing occluded objects, specifically when stimuli become more complicated (e.g. when objects are presented on natural backgrounds).

The additional time needed for processing occluded objects may facilitate object recognition by providing integrated semantic information from visible parts of the target objects, for example, via local recurrent in higher visual areas in the form of attractor networks (Devereux et al., 2018). In other words, partial semantic information (e.g. having wheels, having legs, etc.) may activate prior information associated with the category of the target object (Clarke and Tyler, 2014, Clarke, 2015). Overall these suggest the observed temporal delay under 60% occlusion can be best explained by the engagement of recurrent processes—mostly local recurrent connections.

#### 3.1.2. Computational modeling

Feedforward CNNs have been shown to be able to account for the core object recognition (Cadieu et al., 2014, Khaligh-Razavi and Kriegeskorte, 2014, Yamins et al., 2014, Güçlü and van Gerven, 2015, Kubilius et al., 2016, Kheradpisheh et al., 2016a, Kheradpisheh et al., 2016b, Khaligh-Razavi et al., 2017). The natural question to ask next is whether these models perform similarly well under more challenging conditions, beyond the core object recognition. To address this, we compared a conventional feedforward CNN with a recurrent convolutional network in terms of their object recognition performance, and their representational similarity with that of the human MEG data, under the challenging condition of occlusion. The feedforward CNN only achieved human-level performance when objects were not occluded; and performed significantly lower than the humans and the recurrent network when objects were occluded. The conventional feedforward CNN also failed to explain human neural data when objects were occluded. On the other hand, the convolutional network with local recurrent connections could achieve human-level performance in occluded object recognition and explained a significant variance of the human neural data. Thus, demonstrating that the conventional feedforward CNNs (such as AlexNet) do not account for object recognition under such challenging conditions, where recurrent computations have a prominent contribution.

### 3.2. Unique contribution of recurrent processes in solving object recognition under occlusion

AlexNet (i.e. feedforward model) and HRRN (i.e. recurrent model) both equally explained the early component of the MEG data (<150 ms) with and without occlusion. Consistent with this, the semipartial correlation analyses further revealed no unique variance for these models in explaining the early component of the MEG data. These results suggest that the early component of the MEG data under both conditions (with and without occlusion) are mainly feedforward and both AlexNet and HRRN share a common feedforward component that is significantly correlated with dominantly feedforward MEG representations before 150 ms (Figure S8 shows a plausible Venn diagram describing the relationship between the two models and the MEG data.).

On the other hand, the later component of the MEG data (>150 ms) under occlusion was only correlated with the recurrent model, which had a significant unique contribution in explaining the MEG data under this condition. Under no occlusion, while the later component of the MEG data is significantly correlated with both AlexNet and HRRN, only the HRRN model showed a significant unique contribution in explaining the data (Figure S9). This shows that under no-occlusion the later component of the MEG data can still be partially explained by the common feedforward component of the two models, perhaps because object recognition under no-occlusion is primarily a feedforward process, however the recurrent model has some unique advantages in explaining later MEG components—even under no-occlusion.

### 3.3. Object occlusion vs. object deletion

Object recognition when part of an object is removed without an occluder is one of the challenging conditions that has been previously studied (Nielsen et al., 2006, Wyatte et al., 2012, O’Reilly et al., 2013, Wyatte et al., 2014, Tang et al., 2014, Tang et al., 2018) and may partly look similar to occlusion. However, as shown by Johnson and Olshausen (2005) deleting part of an object is different from occluding it with another object. See Figure S10 for sample images of occlusion and deletion. Occlusion occurs when an object or shape appears in front of another one (Johnson and Olshausen, 2005), in which case the occluding object might serve as an additional cue for object completion. On the other hand, deletion occurs when part of an object is removed without additional cues about the shape or the place of the missing part. Given the difference between these two phenomena at the level of stimulus set, we expect the dynamics of object processing (and the underlying computational mechanisms) to be also different when part of an object is occluded compared to when it is deleted. Consistent with this, Johnson and Olshausen (2005) demonstrated that ERP responses in occipital and parietal electrodes are signifcantly different between object occlusion and deletion. Furthermore, there were significant behavioural differences between object occlusion and deletion, including differneces in recogntion memory and response confidence.

Object deletion has been previously studied in humans using a variety of techniques: Wyatte et. al. (2012, 2014) used human behaviour (in a backward masking paradigm), and computational modelling to show that object recogntion, when part of the object is deleted, requires recurrent processing. Tang et. al. (2014, 2018) used intracranial field potential recording on epileptic patients to study temporal dynamics of object deletion; and proposed an attractor-based recurrent model that could explain the neural data. They found ∼100 ms delay in processing objects when part of the object was deleted, compared to when the whole object was present. In comparison, in our study we found ∼60 ms delay in object processing when objects were occluded. This suggests that while object recognition under both occlusion and deletion requires recurrent processes, temporal dynamics of object deletion is slower, potentially due to the absence of the occluder, which can make the recognition task more difficult.

To summarize, while object deletion has been previously studied in humans, to our knowledge, temporal dynamics of object occlusion had not been studied before. In particular this was the first MVPA study in humans that charachaterized representational dynamics of object recognition under occlusion, and further provided a computational account of the underlying processes that explained both behavioral and neural data.

### 3.4. Does a feedforward system with arbitrarily long depth work the same as a recurrent system with limited depth?

While conventional CNNs could not account for object recognition beyond the core recognition problem, we do not rule out the possibility that much deeper CNNs could perform better under such challenging conditions.

Computational studies have shown that very deep CNNs outperform shallow ones on a variety of object recognition tasks (Simonyan and Zisserman, 2014, Szegedy et al., 2015, Taigman et al., 2014). Specifically, residual learning allows for a much deeper neural network with hundreds (He et al., 2016a) and even thousands (He et al., 2016b) of the layers providing better performance. This is due to the fact that the complex functions that can be represented by deeper architectures cannot be represented by shallow architectures (Bengio and LeCun, 2007). Recent computational modeling studies have tried to clarify why increasing the depth of a network can improve its performance (Liang and Hu, 2015, Liao and Poggio, 2016). These efforts have demonstrated that unfolding a recurrent architecture across time leads to a feedforward network with arbitrary depth, in which the weights are shared among the layers. Although such a recurrent network has far fewer parameters, Liao and Poggio (2016) have empirically shown that it performs as well as a very deep feedforward network *without* shared weights. We also showed that a very deep ResNet (e.g. with 150 layers) can be reformulated into the form of a recurrent CNN with much fewer layers (e.g. five layers) (Figure 6). Thus, a compact architecture that resembles these very deep networks in terms of performance is a recurrent hierarchical network with much fewer layers. This compact architecture is probably what the human visual system has selected to be like (Lamme et al., 1998, Sporns and Zwi, 2004), given the biological constraints of having a very deep neural network inside the brain (Dunbar, 1992, Kaas, 2000, Weaver, 2005, Isler and van Schaik, 2009, Bosman and Aboitiz, 2015).

From a computational viewpoint, recognition of complex images might require more processing efforts; in other words, they might need to go through more layers of processing to be prepared for the final readout. Similarly, in a recurrent architecture, more processing means more iterations. Our modeling results supports this assumption, showing that under more challening recogntion tasks, more iterations are required to reach human-level perfomrance. For example, under 60% and 80% occlusion, the HRRN model reached human level performance, respectively after going through 13 recurrent stages, and 43 recurrent stages (Figure S11). With more iterations, the HRRN model tends to achieve a performance slightly better than the average categorization performance in humans.

Our choice of a recurrent architecture, as opposed to an arbitrarily deep neural network, is mainly driven by the plausibly of such architecture with the hierarchy of vision, where there is only a limited number of processing layers. However, in terms of performance in real-world object recognition tasks (e.g. object recognition under occlusion), the key in achieving a good performance is the number of non-linear operations, which can come either in the form of deeper networks in a feedforward architecture or otherwise more iterations in a recurrent architecture.

### The neural basis of masking effect

Backward masking is a useful tool for studying temporal dynamics of visual object processing (Lamme et al., 2002, Breitmeyer and Öğmen, 2006). It can impair recognition of the target object and reduce or eliminate perceptual visibility through the presentation of a second stimulus (mask) immediately or with an interval after the target stimulus, e.g. 50 ms after the target’s onset. While the origin of masking effect was not the focus of the current study, our MEG results could provide some insights about the underlying mechanisms of backward masking.

There are several accounts of backward masking in the literature: Breitmeyer and Ganz (1976) provided a purely feedforward explanation (two-channel model), arguing that the mask travels rapidly through the fast channel disrupting recognition of the target object traveling through the slow channel. A number of other studies, however, suggest that the masking mainly interferes with the top-down feedback processes (Lamme and Roelfsema, 2000, Lamme et al., 2002, Breitmeyer and Öğmen, 2006, Fahrenfort et al., 2007). And finally, Macknik and Martinez-Conde (2007) explain the masking effect by the lateral inhibition mechanism of neural circuits within different levels of the visual hierarchy; arguing that the mask interferes with the recognition of the target object through lateral inhibition (i.e. inhibitory interactions between target and mask).

The last two accounts of masking, while being different, both argue for the disruption of recurrent processes by the mask: either the top-down recurrent processes, or the local recurrent processes (e.g. lateral interactions). With a short interval between the target and mask, the mask may interfere with the fast recurrent processes (i.e. local recurrent) while with a relatively long interval it may interfere with the slow recurrent processes (i.e. top-down feedback).

Our results of MEG decoding time-courses, time-time decoding and behavioral performances under the no-occlusion condition does not support the purely feedforward account of visual backward masking. We showed that the backward masking did not have a significant effect on disrupting the fast feedforward processes of object recognition under no occlusion (MEG: Figure 4a; behaviorally: Figure 5d). On the other hand, when objects were occluded the backward masking significantly impaired object recognition both behaviorally (Figure 5d) and neurally (Figure 4b). Additionally, the time-time decoding results (Figure 4c, 4d, 4f) showed that backward masking, under no occlusion, had no significant effect on disrupting the diagonal component of the temporal generalization matrix that is mainly associated with the feedforward path of visual processing. On the other hand, the masking removed the off-diagonal components and the late peak (>200ms) observed in the temporal generalization matrix of the occluded objects.

Taken together, our MEG and behavioral results are in favor of a recurrent account for backward masking. Particularly in our experiment with a short stimulus onset asynchrony (SOA = time from stimulus onset to the mask onset), the mask seems to have affected mostly the local recurrent connections.

## 4. Methods

### 4.1. Ethics Statement

The study was conducted according to the Declaration of Helsinki. The experiment protocol was approved by the local committee on the use of humans as experimental subjects. Volunteers completed a consent form before participating in the experiment and were financially compensated after finishing the experiment.

### 4.2. Occluded objects image set

Images of four different object categories (i.e. camel, deer, car, and motorcycle) were used as the stimulus set (Figure 1b). Object images were transformed to be similar in terms of size and contrast level. To generate an occluded image, in an iterative process we added several black circles (as artificial occluders) of different sizes in random positions on the image. The configuration of black circles (i.e. number, size, and their positions on the images) were randomly selected as such that a V1-like model could not discriminate between images with 0%, 60% and 80% occlusion. To simulate the type of occlusion that occurs in natural scenes, the black circles are positioned in both front and back of the target object. The percent of object occlusion is defined as the percent of the target object covered by the black occluders. We defined three levels of occlusion: 0% (no occlusion), 60% occlusion and 80% occlusion. Black circles also existed in the 0% occlusion, but not covering the target object; this was to make sure that the difference observed between occluded and un-occluded objects cannot be solely explained by the presence of these circles. The generated image set is comprised of 12 conditions: four objects × three occlusion levels. For each condition, we generated M = 64 sample images varying by the occlusion pattern and the target object position. To remove the potential effect of low-level visual features in object discrimination—objects positions were slightly changed around the center of the images (by Δx ≤ 15, Δy ≤ 15 pixels). Overall, we controlled for low-level image statistics, as such that images of different levels of occlusion could not be discriminated simply by using low-level visual features (i.e. Gist and V1 model).

### 4.3. Participants and MEG experimental design

Fifteen young volunteers (22-38 year-old, all right-handed; 7 female) participated in the experiment. During the experiment, participants completed eight runs; each run consisted of 192 trials and lasted for approximately eight minutes (total experiment time for each participant = ∼70min). Each trial started with 1sec fixation followed by 34ms (2 × screen frame rate (17ms) = 34ms) presentation of an object image (6° visual angle). In half the trials, we employed backward masking in which a dynamic mask was presented for 102ms shortly after the stimulus offset— inter-stimulus-interval (ISI) of 17ms—(Figure S1). In each run, each object image (i.e. camel, deer, car, motor) was repeated 8 times under different levels of occlusions without backward masking; and another 8 repetitions with backward masking. In other words, each condition (i.e. combination of object-image, occlusion-level, mask or no-mask) was repeated 64 times over the duration of the whole experiment.

Every 1-3 trials, a question mark appeared on the screen (lasted for 1.5 sec) prompting participants to choose animate if the last stimulus was deer/camel and inanimate if the last stimulus was car/motor (Figure S1; see Figure S12 for behavioral performance of animate/inanimate task). Participants were instructed to only respond and blink during the question trials to prevent contamination of MEG signals with motor activity and the eye-blink artifact. The question trials were excluded from further MEG analyses.

The dynamic mask was a sequence of random images (n = 6 images; each presented for 17ms) selected from a pool of the synthesized mask images. They were generated by using a texture synthesis toolbox that is available at: http://www.cns.nyu.edu/~lcv/texture/ (Portilla and Simoncelli, 2000). The synthesized images have low-level feature statistics similar to the original stimuli.

### 4.4. MEG acquisition

To acquire brain signals with millisecond temporal resolution, we used 306-sensors MEG system (Elekta Neuromag, Stockholm). The sampling rate was 1000Hz and band-pass filtered online between 0.03 and 330 Hz. To reduce noise and correct for head movements, raw data were cleaned by spatiotemporal filters [Maxfilter software, Elekta, Stockholm; (Taulu and Simola, 2006)]. Further pre-processing was conducted by Brainstorm toolbox (Tadel et al., 2011). Trials were extracted −200ms to 700ms relative to the stimulus onset. The signals were then normalized by their baseline (−200ms to 0ms), and were temporally smoothed by low-pass filtering at 20Hz.

### 4.5. Behavioral task of multiclass object recognition

We ran a psychophysical experiment, outside MEG, to evaluate human performance on a multi-class occluded object recognition task. Sixteen subjects participated in a session lasting about 40 minutes. The experiment was a combination of mask and no-mask trials that were randomly distributed across the experiment. Each trial, started by a fixation point presented for 0.5s followed by a stimulus presentation of 34ms. In the masked trials, a dynamic mask of 102ms was presented after a short ISI of 17ms (Figure S5). Subjects were instructed to respond accurately and as soon as possible after detecting the target stimulus. They were asked to categorize the object images by pressing one of the pre-assigned four keys on a keyboard corresponding to the four object categories: camel, deer, car, and motorcycle.

Overall, 16 human subjects (25 to 40 years-old) participated in this experiment. Before the experiment, participants performed a short training phase on a totally different image-set to learn the task and reach a predefined performance level in the multi-class object recognition task. The main experiment consisted of 768 trials that were randomly distributed into four blocks of 192 trials (32 repetitions of object images with small variations in position and the pattern of occlusion × three occlusion levels × two masking conditions × four object categories = 768). Images of 256×256 pixels size were presented at a distance of 70 cm at the center of a CRT monitor with the frame rate of 60 Hz and a resolution of 1024×768. We used the MATLAB based psychophysics toolbox of (Pelli, 1997).

### 4.6. Multivariate pattern analyses (MVPA)

#### 4.6.1. Pairwise decoding analysis

To measure temporal dynamics of object information processing, we used pairwise decoding analysis on the MEG data (Isik et al., 2014, Cichy et al., 2014, Kietzmann et al., 2017). For each subject, at each time-point, we created a data matrix of 64-trials × 306-sensors per condition. We used a support vector machine (SVM) to pairwise decode any two conditions, with a leave-one-out cross-validation approach. At each time-point, for each condition, *N-1* pattern vectors were used to train the linear classifier [SVM; LIBSVM, (Chang and Lin, 2011), software available at http://www.csie.ntu.edu.tw/~cjlin/libsvm/], and the remaining *N^th^* vector was used for evaluation. The above procedure was repeated 100 times with random reassignment of the data to training and testing sets. This was then averaged over the six pairwise decoding accuracies. The SVM decoding analysis is done independently for each subject and then we report the average decoding performance over these individuals (Figure 1a).

#### 4.6.2. Time-time decoding analysis

We also reported time-time decoding accuracies, obtained by cross-decoding across time. For each pair of objects, a SVM classifier is trained at a given time and tested at all other time-points, thus showing the generalization of the classifier across time. The results are then averaged across all the pairwise classifications. This yields an 801×801 (t=-100 to 700 ms) matrix of average pairwise decoding accuracies for each subject. Figure 2 shows the time-time decoding matrices averaged across 15 participants. To test for statistical significance, we did one-sided signrank test against the chance-level and then corrected for multiple comparison using FDR.

#### 4.6.3. Sensorwise decoding analysis

We also examined a sensorwise visualization of pairwise object decoding across time (Supp Movies 1 & 2). To this end, we trained and tested the SVM classifier at the level of sensors (i.e. combination of three neighboring sensors) across the whole 306 sensors. First, 306 MEG sensors were grouped into 102 triplets (Elekta Triux system; 2 gradiometers and 1 magnetometer in each location). At each time-point, we applied the same pairwise decoding procedure as previously explained in 4.5.1, this time at the level of groups of 3 adjacent sensors (instead of taking all the 306 MEG sensors together). Average pairwise decoding accuracies across subjects, at each time point, are color-coded across the head surface. We used black dots to indicate channels with significantly above chance accuracy (FDR-corrected across both time and sensors), and gray dots to show accuracies with p<0.05, before correcting for multiple comparison. At each time-point, we also specify the channel with peak decoding accuracy by a red dot.

#### 4.6.4. Representational similarity analysis (RSA) over time

We used representational similarity analysis (RSA) (Kriegeskorte, 2009, Kriegeskorte and Kievit, 2013, Cichy et al., 2014, Carlson et al., 2013, Khaligh-Razavi et al., 2016), to compare representations of computational models with time-resolved representations derived from MEG data.

For the MEG data, representational dissimilarity matrices (RDM) were calculated at each time-point by computing the dissimilarity (1 - Spearman’s R) between all pairs of the MEG patterns elicited by object images. Time-resolved MEG RDMs were then correlated (Spearman’s R) with the computational model RDMs, yielding a correlation vector over time (Figure 5a).

**Semipartial correlation:** We additionally calculated semipartial correlations between the MEG RDMs and the computational model RDMs (Figure 7b). Semipartial correlation indicates the unique relationship between a model representation (e.g. AlexNet RDM) and the MEG data, by taking out the shared contribution of other models (e.g. HRRN RDM) in explaining the MEG data. The semipartial correlation is computed as follows:

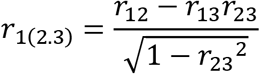

‘*r*_1(2.3)_’ is the correlation between *model X*_1_ and *model X*_2_ controlling for the effect of *model X*_3_ (i.e. removing X_3_ only from X_2_)(Pedzahur, 1997).

To construct CNN model RDMs, we used the extracted features from the penultimate layer of the networks (i.e. the layer before softmax operation). Significant correlations were determined by one-sided signrank test (p < 0.5, FDR-corrected across time).

### 4.7. Significance Testing

We used the non-parametric Wilcoxon signrank test (Gibbons and Chakraborti, 2011) for random effect analysis. To determine time-points with significantly above chance decoding accuracy (or significant RDM correlations), we used a right-sided signrank test across n = 15 participants. To adjust p-values for multiple comparisons (e.g. across time), we further applied the false discovery rate (FDR) correction (Benjamini and Hochberg, 1995) [RSA-Toolbox: is available from https://github.com/rsagroup/rsatoolbox (Nili et al., 2014)].

To determine whether two time-courses (e.g. correlation or decoding) are significantly different at any time interval, we used a two-sided signrank test, FDR corrected across time.

**Onset latency:** We defined onset latency as the earliest time where performance became significantly above chance for at least ten consecutive milliseconds. Mean and standard deviation (SD) for onset latencies were calculated by leave-one-subject-out repeated for N=15 times.

**Peak latency:** The time for peak decoding accuracy was defined as the time where the decoding accuracy was the maximum value. The mean and SD for peak latencies were calculated similar to the onset latencies.

### 4.8. Computational modeling

#### 4.8.1. Feedforward computational model (AlexNet)

We used a well-known CNN (AlexNet) (Krizhevsky et al., 2012) that is shown to account for the core object recognition (Khaligh-Razavi and Kriegeskorte, 2014, Cadieu et al., 2014, Kheradpisheh et al., 2016a, Kheradpisheh et al., 2016b). CNNs are cascades of hierarchically organized feature extraction layers. Each layer has several hundred convolutional filters and each convolutional filter scans various places on the input generating a feature map at its output. A convolutional layer may be followed by a local or global pooling layer merging outputs of a group of units. The pooling layers make the feature maps invariant to small variations (Bengio and LeCun, 2007). AlexNet has eight cascading layers: five convolutional layers, some of which followed by pooling layers, and three fully-connected layers (Krizhevsky et al., 2012). The last fully-connected layer is a 1000-way softmax that corresponds to the 1000 class labels. The network has 60 million free parameters. A pre-trained version of the model, which is trained on 1.2 million images from ImageNet dataset (Russakovsky et al., 2015) is used for the experiments here. We used the features extracted by the fc7 layer (before softmax operation) as the model output.

#### 4.8.2. Hierarchical Recurrent ResNet (HRRN)

In convolutional neural networks, performance in visual recognition tasks can be substantially improved by adding to the depth of the network (Simonyan and Zisserman, 2014, Szegedy et al., 2015, He et al., 2015). However, this comes at a cost: deeper networks of simply stacking layers (plain nets) have higher training errors due to the vanishing gradients (degradation) (Glorot and Bengio, 2010) problem that prevents convergence in the training phase. To address this problem, He et al. (2016a) introduced a deep residual learning framework. Residual networks can overcome the vanishing gradient problem during learning by employing *identity shortcut connections* that allow bypassing residual layers. This framework enables training ultra-deep networks, e.g. with 1202 layers, leading to much better performances compared to the shallower networks (He et al., 2016a, He et al., 2016b).

Residual connections give ResNet an interesting characteristic of having several possible pathways with different lengths from the network’s input to the output instead of a single deep pathway (Veit et al., 2016). For example, the ultra-deep 152-layers ResNet in its simplest form—by skipping all the residual layers—is a hierarchy of five convolutional layers. By including additional residual layers, more complex networks with various depths are constructed [see table 1 in (He et al., 2016a)]. Each series of the residual modules can be reformulated into the form of a convolutional layer with recurrent connections (Liang and Hu, 2015). Additionally, Liao and Poggio (2015) show that a ResNet with shared weights can retain most of the performance of the corresponding network with non-shared weights.

In this study, we proposed a generalization of this convolutional neural network by redefining residual layers as local recurrent connections. As shown in Figure 5, we reformulated the 152- layers ResNet of He et al. (2016a) into the form of a five-layer convolutional network with folded residual layers as its local recurrent connections. The unfolded HRRN (=152 layers ResNet) is deeper than the AlexNet and has different normalization (i.e. batch normalization) and filter sizes, however, they still have a similar number of free parameters (60M). Comparing AlexNet with a purely feedforward version of HRRN (readout stage 0, with five layers), AlexNet performs slightly better than HRRN in readout stage 0, and gradually after 4 iterations HRRN reaches a categorization performance similar to that of AlexNet (Figure S13). The model is pre-trained on ImageNet 2012 dataset with a training set similar to that of Alexnet (1.2 million training images).

It is shown experimentally that an unfolded recurrent CNN (with shared weights) is similar to a very deep feedforward network with non-shared weights (Liao and Poggio, 2016). In our analyses, we used the extracted features of the penultimate layer (i.e. layer pool5, which is before the softmax layer) as the model output.

## Acknowledgement

The study was conducted at the Athinoula A. Martinos Imaging Center at the McGovern Institute for Brain Research, Massachusetts Institute of Technology. We would like to thank Aude Oliva and Dimitrios Pantazis for their help and support in conducting this study. We would also like to thank Radoslaw Martin Cichy for helpful comments. SKR was funded by a return home fellowship grant from Iranian national elite foundation.

## References

Bacon-Macé, N., Macé, M. J.-M., Fabre-Thorpe, M. & Thorpe, S. J. 2005. The time course of visual processing: Backward masking and natural scene categorisation. Vision research, 45, 1459–1469.

Ban, H., Yamamoto, H., Hanakawa, T., Urayama, S.-I., Aso, T., Fukuyama, H. & Ejima, Y. 2013. Topographic representation of an occluded object and the effects of spatiotemporal context in human early visual areas. Journal of Neuroscience, 33, 16992–17007.

Bengio, Y. & Lecun, Y. 2007. Scaling learning algorithms towards AI. Large-scale kernel machines, 34.

Benjamini, Y. & Hochberg, Y. 1995. Controlling the false discovery rate: a practical and powerful approach to multiple testing. Journal of the royal statistical society. Series B (Methodological), 289–300.

Bosman, C. A. & Aboitiz, F. 2015. Functional constraints in the evolution of brain circuits. Frontiers in neuroscience, 9.

Breitmeyer, B. & Öğmen, H. 2006. Visual masking: Time slices through conscious and unconscious vision, Oxford University Press.

Cadieu, C. F., Hong, H., Yamins, D. L., Pinto, N., Ardila, D., Solomon, E. A., Majaj, N. J. & Dicarlo, J. J. 2014. Deep neural networks rival the representation of primate IT cortex for core visual object recognition. PLoS Comput Biol, 10, e1003963.

Carlson, T., Tovar, D. A., Alink, A. & Kriegeskorte, N. 2013. Representational dynamics of object vision: the first 1000 ms. Journal of vision, 13, 1–1.

Chang, C.-C. & Lin, C.-J. 2011. LIBSVM: a library for support vector machines. ACM transactions on intelligent systems and technology (TIST), 2, 27.

Choi, H., Pasupathy, A. & Shea-Brown, E. 2016. Predictive coding in area V4: dynamic shape discrimination under partial occlusion. arXiv preprint arXiv:1612.05321.

Cichy, R. M., Khosla, A., Pantazis, D., Torralba, A. & Oliva, A. 2016a. Comparison of deep neural networks to spatio-temporal cortical dynamics of human visual object recognition reveals hierarchical correspondence. Scientific reports, 6, 27755.

Cichy, R. M., Pantazis, D. & Oliva, A. 2014. Resolving human object recognition in space and time. Nature neuroscience, 17, 455–462.

Cichy, R. M., Pantazis, D. & Oliva, A. 2016b. Similarity-based fusion of MEG and fMRI reveals spatio-temporal dynamics in human cortex during visual object recognition. Cerebral Cortex, 26, 3563–3579.

Clarke, A. 2015. Dynamic information processing states revealed through neurocognitive models of object semantics. Language, cognition and neuroscience, 30, 409–419.

Clarke, A., Taylor, K. I. & Tyler, L. K. 2011. The evolution of meaning: spatio-temporal dynamics of visual object recognition. Journal of cognitive neuroscience, 23, 1887–1899.

Clarke, A. & Tyler, L. K. 2014. Object-specific semantic coding in human perirhinal cortex. Journal of Neuroscience, 34, 4766–4775.

Clarke, A. M., Herzog, M. H. & Francis, G. 2014. Visual crowding illustrates the inadequacy of local vs. global and feedforward vs. feedback distinctions in modeling visual perception. Frontiers in psychology, 5.

Contini, E. W., Wardle, S. G. & Carlson, T. A. 2017. Decoding the time-course of object recognition in the human brain: From visual features to categorical decisions. Neuropsychologia.

Deng, J., Dong, W., Socher, R., Li, L.-J., Li, K. & Fei-Fei, L. Imagenet: A large-scale hierarchical image database. Computer Vision and Pattern Recognition, 2009. CVPR 2009. IEEE Conference on, 2009. IEEE, 248–255.

Devereux, B. J., Clarke, A. D. & Tyler, L. K. 2018. Integrated deep visual and semantic attractor neural networks predict fMRI pattern-information along the ventral object processing pathway. Scientific Reports.

Dicarlo, J. J. & Cox, D. D. 2007. Untangling invariant object recognition. Trends in cognitive sciences, 11, 333–341.

Dicarlo, J. J., Zoccolan, D. & Rust, N. C. 2012. How does the brain solve visual object recognition? Neuron, 73, 415–434.

Dunbar, R. I. 1992. Neocortex size as a constraint on group size in primates. Journal of human evolution, 22, 469–493.

Eberhardt, S., Cader, J. G. & Serre, T. How deep is the feature analysis underlying rapid visual categorization? Advances in neural information processing systems, 2016. 1100–1108.

Erlikhman, G. & Caplovitz, G. P. 2017. Decoding information about dynamically occluded objects in visual cortex. NeuroImage, 146, 778–788.

Fahrenfort, J. J., Scholte, H. S. & Lamme, V. A. 2007. Masking disrupts reentrant processing in human visual cortex. Journal of cognitive neuroscience, 19, 1488–1497.

Felleman, D. J. & Van Essen, D. C. 1991. Distributed hierarchical processing in the primate cerebral cortex. Cerebral cortex, 1, 1–47.

Garrido, M. I., Kilner, J. M., Kiebel, S. J. & Friston, K. J. 2007. Evoked brain responses are generated by feedback loops. Proceedings of the National Academy of Sciences, 104, 20961–20966.

Ghodrati, M., Farzmahdi, A., Rajaei, K., Ebrahimpour, R. & Khaligh-Razavi, S.-M. 2014. Feedforward object-vision models only tolerate small image variations compared to human. Frontiers in computational neuroscience, 8, 74.

Gibbons, J. D. & Chakraborti, S. 2011. Nonparametric statistical inference. International encyclopedia of statistical science. Springer.

Gilbert, C. D. & Li, W. 2013. Top-down influences on visual processing. Nature Reviews Neuroscience, 14, 350–363.

Glorot, X. & Bengio, Y. Understanding the difficulty of training deep feedforward neural networks. Proceedings of the Thirteenth International Conference on Artificial Intelligence and Statistics, 2010. 249–256.

Goddard, E., Carlson, T. A., Dermody, N. & Woolgar, A. 2016. Representational dynamics of object recognition: Feedforward and feedback information flows. NeuroImage, 128, 385–397.

Grootswagers, T. & Carlson, T. 2015. Decoding the emerging representation of degraded visual objects in the human brain. Journal of vision, 15, 1087–1087.

Grootswagers, T., Wardle, S. G. & Carlson, T. A. 2017. Decoding dynamic brain patterns from evoked responses: A tutorial on multivariate pattern analysis applied to time series neuroimaging data. Journal of cognitive neuroscience.

Güçlü, U. & Van Gerven, M. A. 2015. Deep neural networks reveal a gradient in the complexity of neural representations across the ventral stream. Journal of Neuroscience, 35, 10005–10014.

He, K., Zhang, X., Ren, S. & Sun, J. Delving deep into rectifiers: Surpassing human-level performance on imagenet classification. Proceedings of the IEEE international conference on computer vision, 2015. 1026–1034.

He, K., Zhang, X., Ren, S. & Sun, J. Deep residual learning for image recognition. Proceedings of the IEEE conference on computer vision and pattern recognition, 2016a. 770–778.

He, K., Zhang, X., Ren, S. & Sun, J. Identity mappings in deep residual networks. European Conference on Computer Vision, 2016b. Springer, 630–645.

Hegdé, J., Fang, F., Murray, S. O. & Kersten, D. 2008. Preferential responses to occluded objects in the human visual cortex. Journal of vision, 8, 16–16.

Hulme, O. J. & Zeki, S. 2007. The sightless view: neural correlates of occluded objects. Cerebral Cortex, 17, 1197–1205.

Isik, L., Meyers, E. M., Leibo, J. Z. & Poggio, T. 2014. The dynamics of invariant object recognition in the human visual system. Journal of neurophysiology, 111, 91–102.

Isler, K. & Van Schaik, C. P. 2009. The expensive brain: a framework for explaining evolutionary changes in brain size. Journal of Human Evolution, 57, 392–400.

Johnson, J. S. & Olshausen, B. A. 2005. The recognition of partially visible natural objects in the presence and absence of their occluders. Vision research, 45, 3262–3276.

Kaas, J. H. 2000. Why is brain size so important: Design problems and solutions as neocortex gets biggeror smaller. Brain and Mind, 1, 7–23.

Kafaligonul, H., Breitmeyer, B. G. & Öğmen, H. 2015. Feedforward and feedback processes in vision. Frontiers in psychology, 6.

Kaneshiro, B., Guimaraes, M. P., Kim, H.-S., Norcia, A. M. & Suppes, P. 2015. A Representational Similarity Analysis of the Dynamics of Object Processing Using Single-Trial EEG Classification. PloS one, 10, e0135697.

Karimi-Rouzbahani, H., Bagheri, N. & Ebrahimpour, R. 2017. Hard-wired feed-forward visual mechanisms of the brain compensate for affine variations in object recognition. Neuroscience, 349, 48–63.

Khaligh-Razavi, S.-M., Bainbridge, W. A., Pantazis, D. & Oliva, A. 2016. From what we perceive to what we remember: Characterizing representational dynamics of visual memorability. bioRxiv, 049700.

Khaligh-Razavi, S.-M., Carlin, J., Martin, C. R. & Kriegeskorte, N. 2015. The effects of recurrent dynamics on ventral-stream representational geometry. Journal of vision, 15, 1089–1089.

Khaligh-Razavi, S.-M., Henriksson, L., Kay, K. & Kriegeskorte, N. 2017. Fixed versus mixed RSA: Explaining visual representations by fixed and mixed feature sets from shallow and deep computational models. Journal of Mathematical Psychology, 76, 184–197.

Khaligh-Razavi, S.-M. & Kriegeskorte, N. 2014. Deep supervised, but not unsupervised, models may explain IT cortical representation. PLoS Comput Biol, 10, e1003915.

Kheradpisheh, S. R., Ghodrati, M., Ganjtabesh, M. & Masquelier, T. 2016a. Deep networks can resemble human feed-forward vision in invariant object recognition. Scientific reports, 6, 32672.

Kheradpisheh, S. R., Ghodrati, M., Ganjtabesh, M. & Masquelier, T. 2016b. Humans and deep networks largely agree on which kinds of variation make object recognition harder. Frontiers in computational neuroscience, 10.

Kietzmann, T. C., Gert, A. L., Tong, F. & König, P. 2017. Representational dynamics of facial viewpoint encoding. Journal of cognitive neuroscience, 29, 637–651.

King, J.-R., Pescetelli, N. & Dehaene, S. 2016. Brain mechanisms underlying the brief maintenance of seen and unseen sensory information. Neuron, 92, 1122–1134.

King, J. & Dehaene, S. 2014. Characterizing the dynamics of mental representations: the temporal generalization method. Trends in cognitive sciences, 18, 203–210.

Klink, P. C., Dagnino, B., Gariel-Mathis, M.-A. & Roelfsema, P. R. 2017. Distinct Feedforward and Feedback Effects of Microstimulation in Visual Cortex Reveal Neural Mechanisms of Texture Segregation. Neuron.

Kosai, Y., El-Shamayleh, Y., Fyall, A. M. & Pasupathy, A. 2014. The role of visual area V4 in the discrimination of partially occluded shapes. Journal of Neuroscience, 34, 8570–8584.

Kriegeskorte, N. 2009. Relating population-code representations between man, monkey, and computational models. Frontiers in Neuroscience, 3, 35.

Kriegeskorte, N. & Kievit, R. A. 2013. Representational geometry: integrating cognition, computation, and the brain. Trends in cognitive sciences, 17, 401–412.

Krizhevsky, A., Sutskever, I. & Hinton, G. E. Imagenet classification with deep convolutional neural networks. Advances in neural information processing systems, 2012. 1097–1105.

Kubilius, J., Bracci, S. & De Beeck, H. P. O. 2016. Deep neural networks as a computational model for human shape sensitivity. PLoS computational biology, 12, e1004896.

Lamme, V. A. & Roelfsema, P. R. 2000. The distinct modes of vision offered by feedforward and recurrent processing. Trends in neurosciences, 23, 571–579.

Lamme, V. A., Super, H. & Spekreijse, H. 1998. Feedforward, horizontal, and feedback processing in the visual cortex. Current opinion in neurobiology, 8, 529–535.

Lamme, V. A., Zipser, K. & Spekreijse, H. 2002. Masking interrupts figure-ground signals in V1. Journal of cognitive neuroscience, 14, 1044–1053.

Liang, M. & Hu, X. Recurrent convolutional neural network for object recognition. Proceedings of the IEEE Conference on Computer Vision and Pattern Recognition, 2015. 3367–3375.

Liao, Q. & Poggio, T. 2016. Bridging the gaps between residual learning, recurrent neural networks and visual cortex. arXiv preprint arXiv:1604.03640.

Liu, H., Agam, Y., Madsen, J. R. & Kreiman, G. 2009. Timing, timing, timing: fast decoding of object information from intracranial field potentials in human visual cortex. Neuron, 62, 281–290.

Livne, T. & Sagi, D. 2011. Multiple levels of orientation anisotropy in crowding with Gabor flankers. Journal of vision, 11, 18–18.

Macknik, S. L. & Martinez-Conde, S. 2007. The role of feedback in visual masking and visual processing. Advances in cognitive psychology, 3, 125–152.

Manassi, M. & Herzog, M. Crowding and grouping: how much time is needed to process good Gestalt? Perception, 2013. 229.

Mohsenzadeh, Y., Qin, S., Cichy, R. M. & Pantazis, D. 2018. Ultra-Rapid serial visual presentation reveals dynamics of feedforward and feedback processes in the ventral visual pathway. Elife, 7, e36329.

Nielsen, K. J., Logothetis, N. K. & Rainer, G. 2006. Dissociation between local field potentials and spiking activity in macaque inferior temporal cortex reveals diagnosticity-based encoding of complex objects. Journal of Neuroscience, 26, 9639–9645.

Nili, H., Wingfield, C., Walther, A., Su, L., Marslen-Wilson, W. & Kriegeskorte, N. 2014. A toolbox for representational similarity analysis. PLoS Comput Biol, 10, e1003553.

O’reilly, R. C., Wyatte, D., Herd, S., Mingus, B. & Jilk, D. J. 2013. Recurrent processing during object recognition. Frontiers in psychology, 4, 124.

Pedzahur, E. 1997. Multiple regression in behavioral research: Explanation and prediction. London, UK: Wadsworth, Thompson Learning.

Pelli, D. G. 1997. The VideoToolbox software for visual psychophysics: Transforming numbers into movies. Spatial vision, 10, 437–442.

Portilla, J. & Simoncelli, E. P. 2000. A parametric texture model based on joint statistics of complex wavelet coefficients. International journal of computer vision, 40, 49–70.

Rajalingham, R., Issa, E. B., Bashivan, P., Kar, K., Schmidt, K. & Dicarlo, J. J. 2018. Large-scale, high-resolution comparison of the core visual object recognition behavior of humans, monkeys, and state-of-the-art deep artificial neural networks. bioRxiv, 240614.

Rauschenberger, R., Liu, T., Slotnick, S. D. & Yantis, S. 2006. Temporally unfolding neural representation of pictorial occlusion. Psychological Science, 17, 358–364.

Rensink, R. A. & Enns, J. T. 1998. Early completion of occluded objects. Vision research, 38, 2489–2505.

Russakovsky, O., Deng, J., Su, H., Krause, J., Satheesh, S., Ma, S., Huang, Z., Karpathy, A., Khosla, A. & Bernstein, M. 2015. Imagenet large scale visual recognition challenge. International Journal of Computer Vision, 115, 211–252.

Serre, T., Oliva, A. & Poggio, T. 2007. A feedforward architecture accounts for rapid categorization. Proceedings of the National Academy of Sciences, 104, 6424–6429.

Simonyan, K. & Zisserman, A. 2014. Very deep convolutional networks for large-scale image recognition. arXiv preprint arXiv:1409.1556.

Spoerer, C., Mcclure, P. & Kriegeskorte, N. 2017. Recurrent Convolutional Neural Networks: A Better Model Of Biological Object Recognition Under Occlusion. bioRxiv, 133330.

Sporns, O. & Zwi, J. D. 2004. The small world of the cerebral cortex. Neuroinformatics, 2, 145–162.

Szegedy, C., Liu, W., Jia, Y., Sermanet, P., Reed, S., Anguelov, D., Erhan, D., Vanhoucke, V. & Rabinovich, A. Going deeper with convolutions. Proceedings of the IEEE Conference on Computer Vision and Pattern Recognition, 2015. 1–9.

Tadel, F., Baillet, S., Mosher, J. C., Pantazis, D. & Leahy, R. M. 2011. Brainstorm: a user-friendly application for MEG/EEG analysis. Computational intelligence and neuroscience, 2011, 8.

Taigman, Y., Yang, M., Ranzato, M. A. & Wolf, L. Deepface: Closing the gap to human-level performance in face verification. Proceedings of the IEEE Conference on Computer Vision and Pattern Recognition, 2014. 1701–1708.

Tang, H., Buia, C., Madhavan, R., Crone, N. E., Madsen, J. R., Anderson, W. S. & Kreiman, G. 2014. Spatiotemporal dynamics underlying object completion in human ventral visual cortex. Neuron, 83, 736–748.

Tang, H., Schrimpf, M., Lotter, W., Moerman, C., Paredes, A., Caro, J. O., Hardesty, W., Cox, D. & Kreiman, G. 2018. Recurrent computations for visual pattern completion. Proceedings of the National Academy of Sciences, 201719397.

Taulu, S. & Simola, J. 2006. Spatiotemporal signal space separation method for rejecting nearby interference in MEG measurements. Physics in Medicine & Biology, 51, 1759.

Thorpe, S. J. 2009. The speed of categorization in the human visual system. Neuron, 62, 168–170.

Thorpe, S. J. & Fabre-Thorpe, M. 2001. Seeking categories in the brain. Science, 291, 260–263.

Tomita, H., Ohbayashi, M., Nakahara, K., Hasegawa, I. & Miyashita, Y. 1999. Top-down signal from prefrontal cortex in executive control of memory retrieval. Nature, 401, 699.

Veit, A., Wilber, M. J. & Belongie, S. Residual networks behave like ensembles of relatively shallow networks. Advances in Neural Information Processing Systems, 2016. 550–558.

Weaver, A. H. 2005. Reciprocal evolution of the cerebellum and neocortex in fossil humans. Proceedings of the National Academy of Sciences of the United States of America, 102, 3576–3580.

Wen, H., Shi, J., Chen, W. & Liu, Z. 2018. Deep Residual Network Predicts Cortical Representation and Organization of Visual Features for Rapid Categorization. Scientific Reports, 8, 3752.

Wyatte, D., Curran, T. & O’reilly, R. 2012. The limits of feedforward vision: Recurrent processing promotes robust object recognition when objects are degraded. Journal of Cognitive Neuroscience, 24, 2248–2261.

Wyatte, D., Jilk, D. J. & O’reilly, R. C. 2014. Early recurrent feedback facilitates visual object recognition under challenging conditions.

Yamins, D. L., Hong, H., Cadieu, C. F., Solomon, E. A., Seibert, D. & Dicarlo, J. J. 2014. Performance-optimized hierarchical models predict neural responses in higher visual cortex. Proceedings of the National Academy of Sciences, 111, 8619–8624.

