## Supporting information for "Beyond Core Object Recognition: Recurrent processes account for object recognition under occlusion"

### Supplementary Information

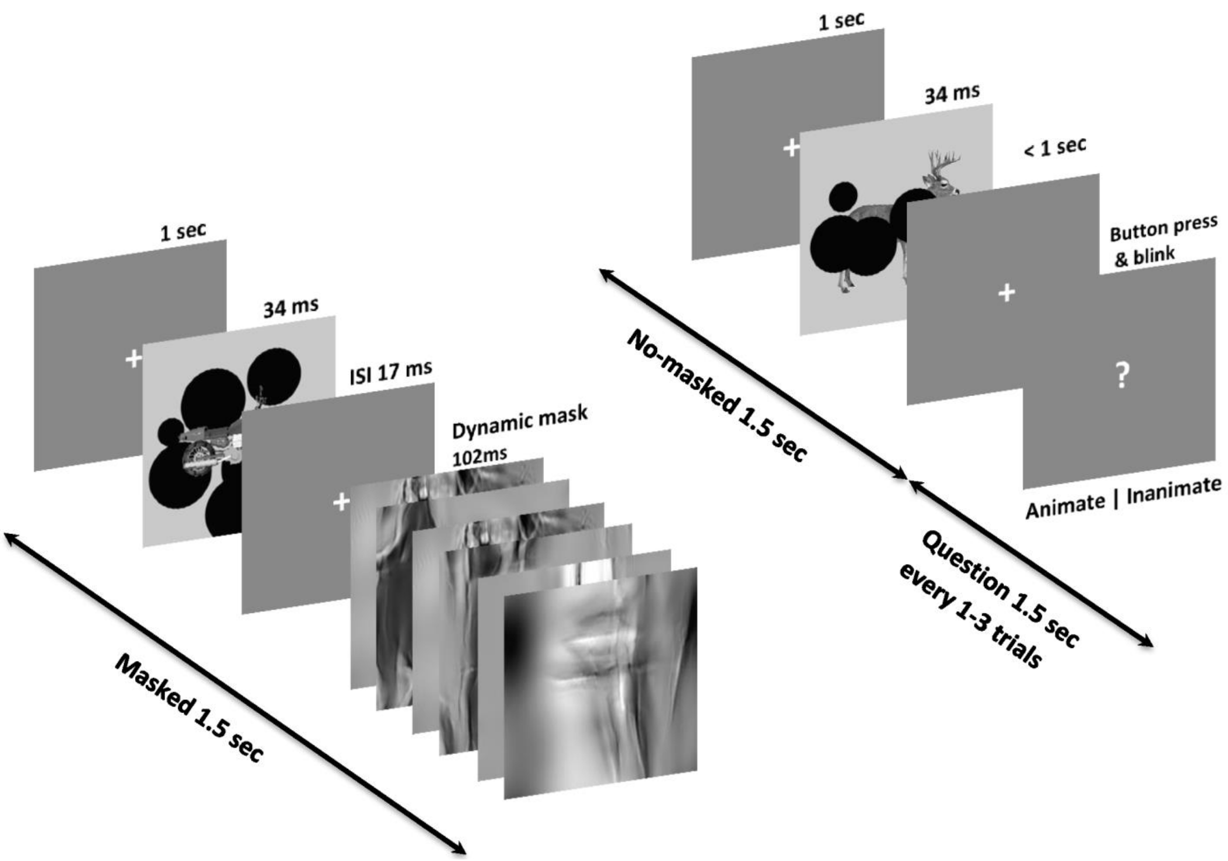


**Figure S1. MEG experimental paradigm.** The experiment was divided into two types of trials: mask and no mask trials, shown in random order. Each trial started by a fixation of 1 sec followed by a target stimulus presented for 34ms. In the mask trials, after a short inter-stimulus-interval (ISI) of 17ms, a dynamic mask of 102ms duration was presented. Every 1-3 trials (average = 2) a question mark appeared on the screen. Subjects were asked to select whether the last image was animate or inanimate. They were also instructed to restrict their blinking (and swallowing) to the question-mark trials.

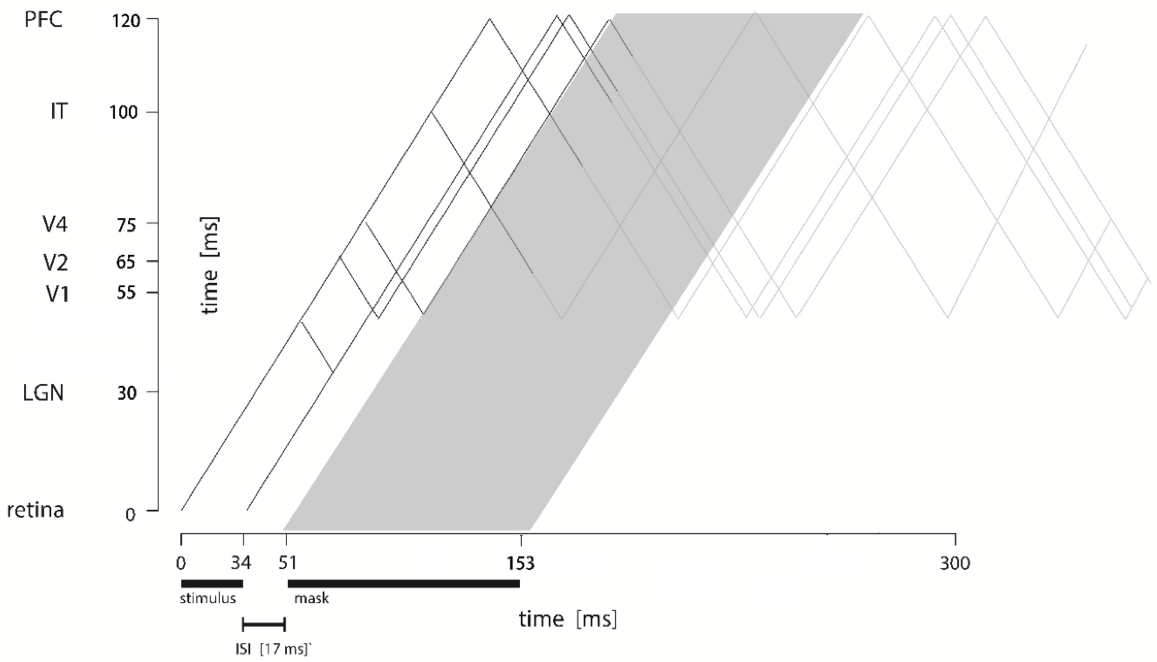


**Figure S2. Time course of visual processing and masking in humans.** Earliest responses reaching each of the visual areas from V1 to IT are indicated by the oblique lines, when a stimulus is on for 34ms, followed by an ISI of 17 ms, followed by a mask. The grey shaded area indicates the effect of mask when it disrupts the information that is being fed back from higher visual areas to lower visual areas. The approximate timings are set according to human (Mormann et al., 2008, Liu et al., 2009, Cichy et al., 2016b) and non-human (i.e. macaque) studies (Lamme and Roelfsema, 2000) controlling for the fact that the macaque cortex is smaller, with a shorter neural distance and therefore faster transmission of visual information (Thorpe and Fabre-Thorpe, 2001).

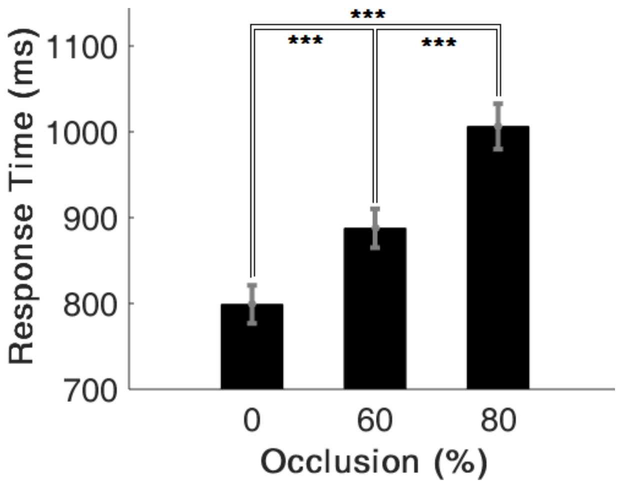


**Figure S3. Average response times of behavioral experiment across three occlusion levels.** The results are averaged over n=15 human participants. Error bars represent SEM. Significant difference between occlusion levels are indicated by stars (signrank test). *** = $p<0.001$.

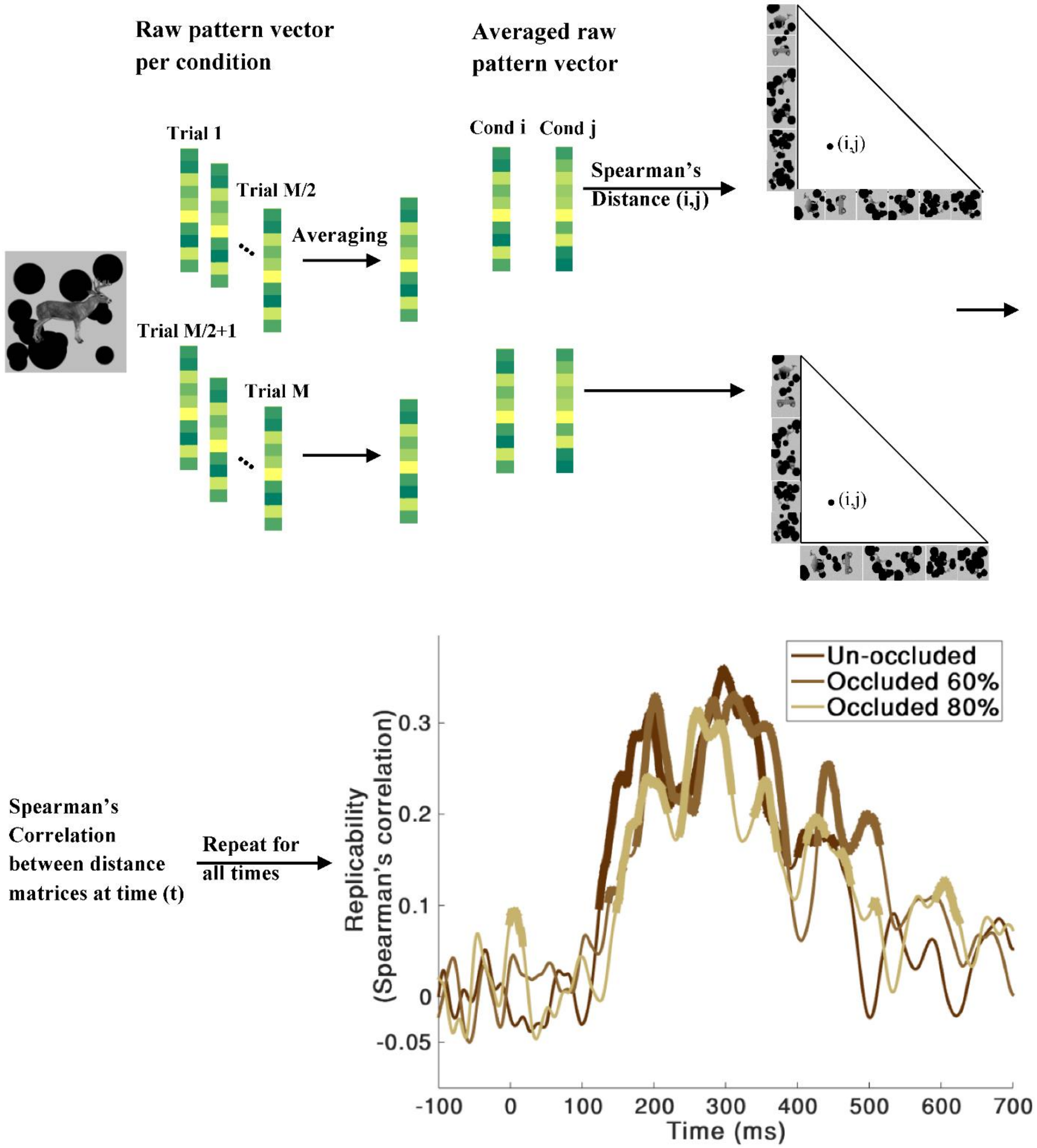


**Figure S4. Split-half replicability for different conditions.** The MEG trials for each condition (i.e. 0% occlusion, 60% occlusion, and 80% occlusion) were divided into two halves, the replicability is measured as the correlation between these two halves. Thicker lines indicate significantly above chance correlations (right sided sign-rank test, FDR corrected across time, $p<0.05$). No significant difference was observed between the replicability of different conditions (two-sided signrank test, FDR-corrected across time), thus indicating that different conditions do not differ in their level of noise. In more details, for each condition, we randomly split M = 64 trial repetitions into two groups of 32 trials. Distance matrices were then calculated for average raw pattern vectors of each group by computing pairwise dissimilarity (1-correlation) between the patterns (12x12 matrices; 12 experimental stimuli). Spearman's R was used as the replicability measure between these two split-half matrices across time.

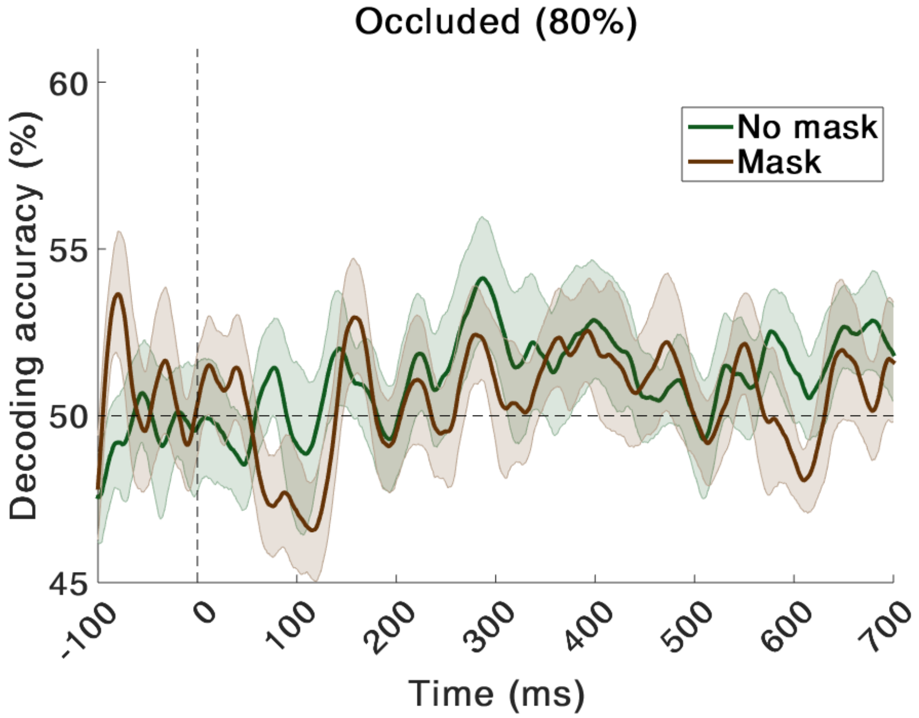


**Figure S5. Time-course of average pairwise decoding accuracy for mask and no mask trials under 80 percent occlusion.** Shaded error bars represent standard error of the mean (SEM). Decoding accuracy was not significantly above chance at any time-point for both mask and no mask (right-sided signrank test, FDR-corrected across time, $p < 0.5$).

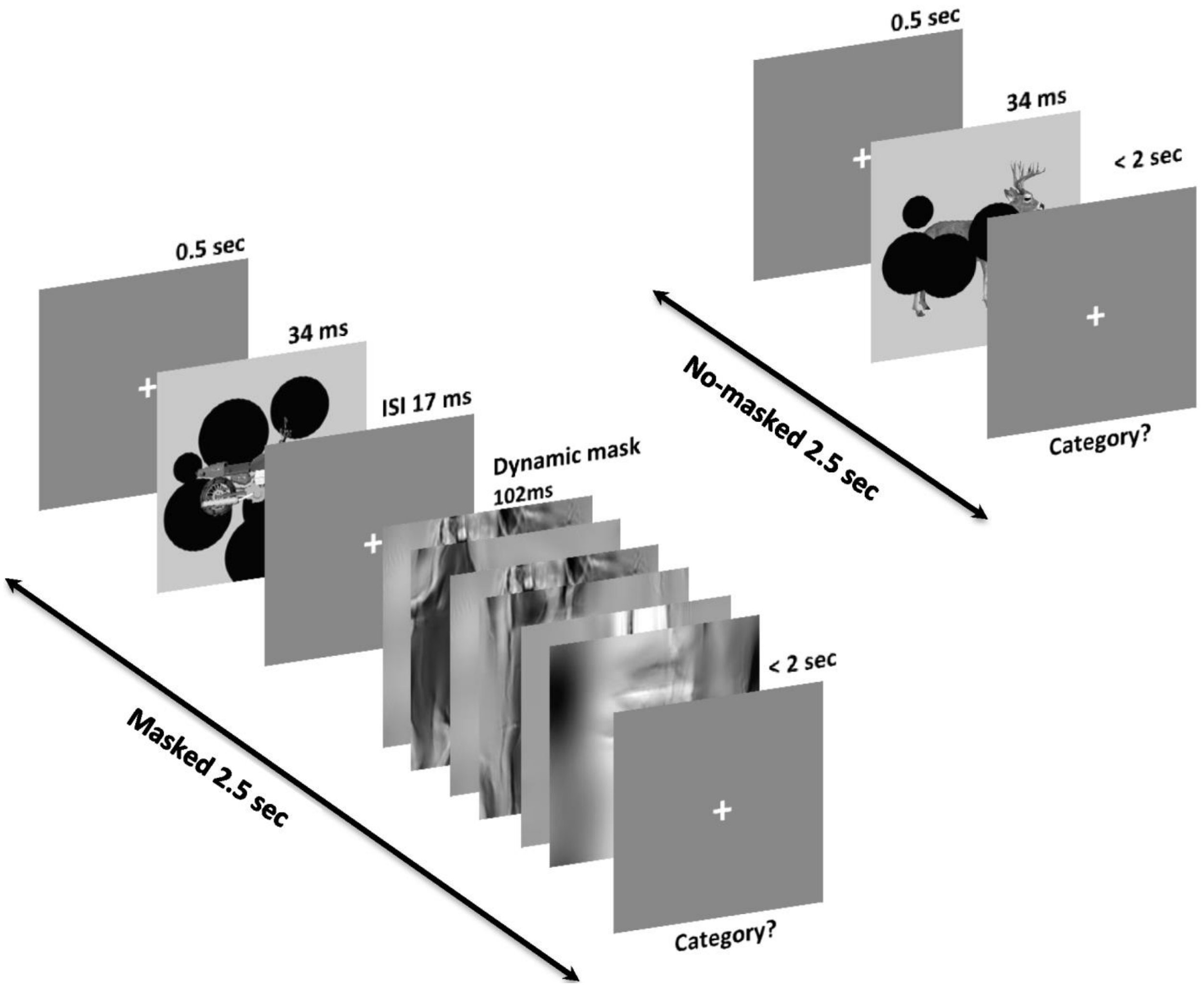


**Figure S6. Experimental design of the multiclass behavioral task.** The behavioral experiment had two types of trials: mask and no mask trials (in random order). Each trial started by 0.5sec fixation, followed by a short presentation of stimulus for 34ms. In the masked trials, 17ms after the stimulus offset (short ISI) a dynamic mask of 100ms was presented. The dynamic mask was a sequence of synthesized images. The subjects were instructed to respond as soon and accurate as possible. Subject's response was to categorize the presented image by pressing one of the four pre-specified keys on a keyboard corresponding to the four object categories (camel, deer, car, and motorcycle).

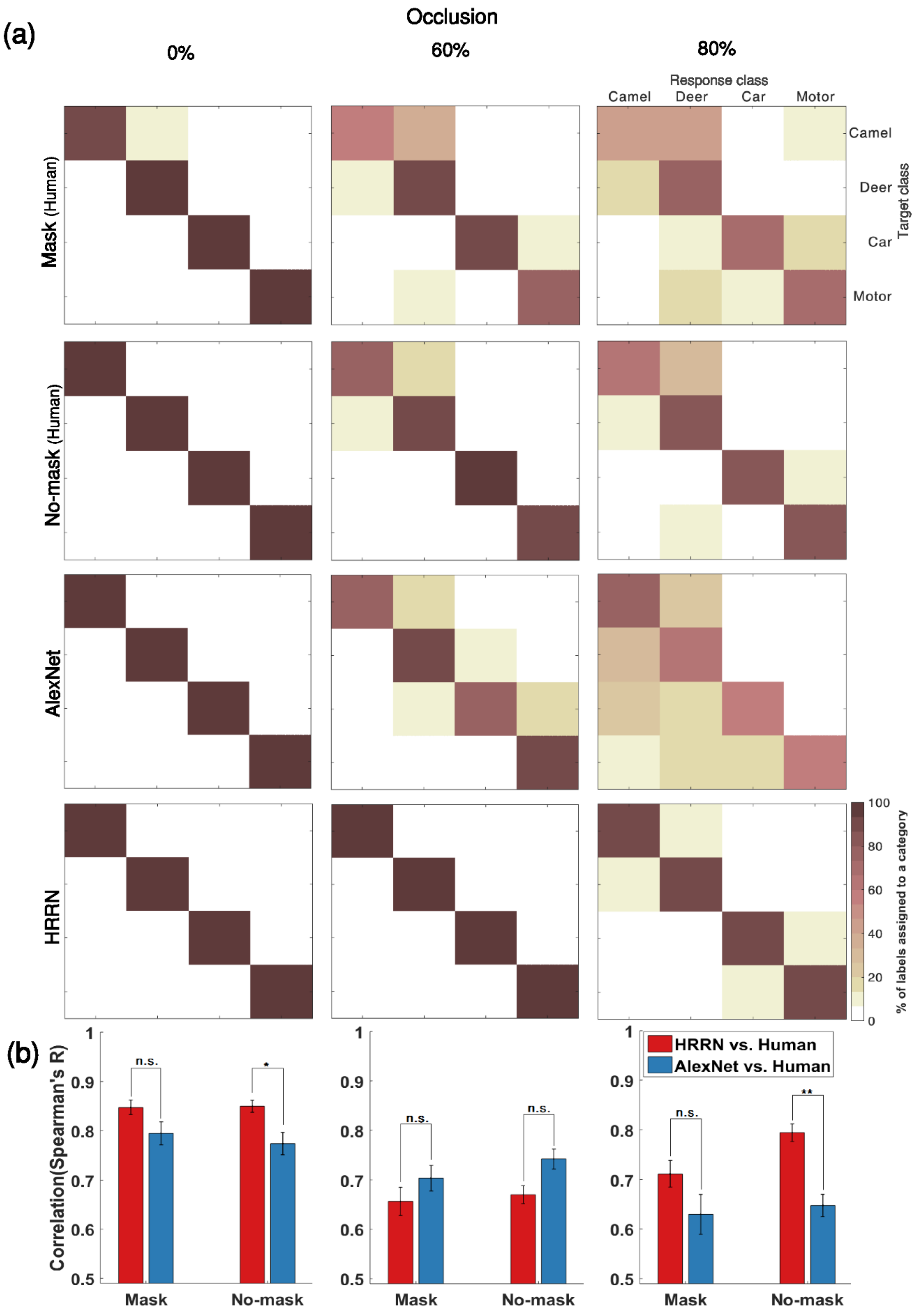
(a)
Occlusion
0%
60%
80%
Response class
Camel
Deer
Car
Motor
Target class
Camel
Deer
Car
Motor
Mask (Human)
No-mask (Human)
AlexNet
HRRN
% of labels assigned to a category
100
80
60
40
20
0
(b)
Correlation(Spearman's R)
1
0.9
0.8
0.7
0.6
0.5
n.s.
*
**
Mask
No-mask
HRRN vs. Human
AlexNet vs. Human

69

**Figure S7. Confusion matrices of the human (mask/no-mask) and models across three occlusion levels.** To compare patterns of errors in the models and humans, we computed confusion matrices. To obtain a confusion matrix, we first trained a SVM classifier on a multiclass object recognition task similar to the behavioral experiment. Then, we calculated the percentage of predicted labels assigned to a category. We display these percentages using color-codes in the matrix. Elements in the main diagonal of the confusion matrix show classification performances and off-diagonal elements show errors made in the classification. **(a)** Confusion matrices for the three levels of occlusion. The color bar, at the bottom-right corner, indicates the percentage of labels assigned to a category. **(b)** Bars indicate correlations between confusion matrices of the models with that of humans (mask and no-mask). Stars show significant differences between HRRN and AlexNet (signrank test, across subjects). * = $p<0.05$; ** = $p<0.01$.

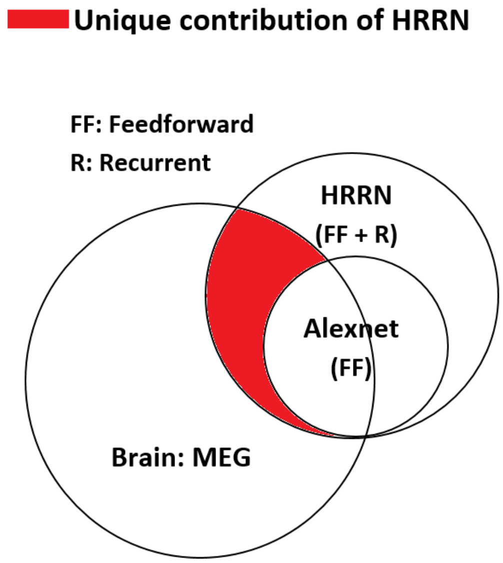


**Figure S8. Venn diagram of MEG and the models.** Red area indicates the unique contribution of HRRN in explaining MEG data. AlexNet has no unique contribution likely due to a component shared between the two models (i.e. feedforward component).

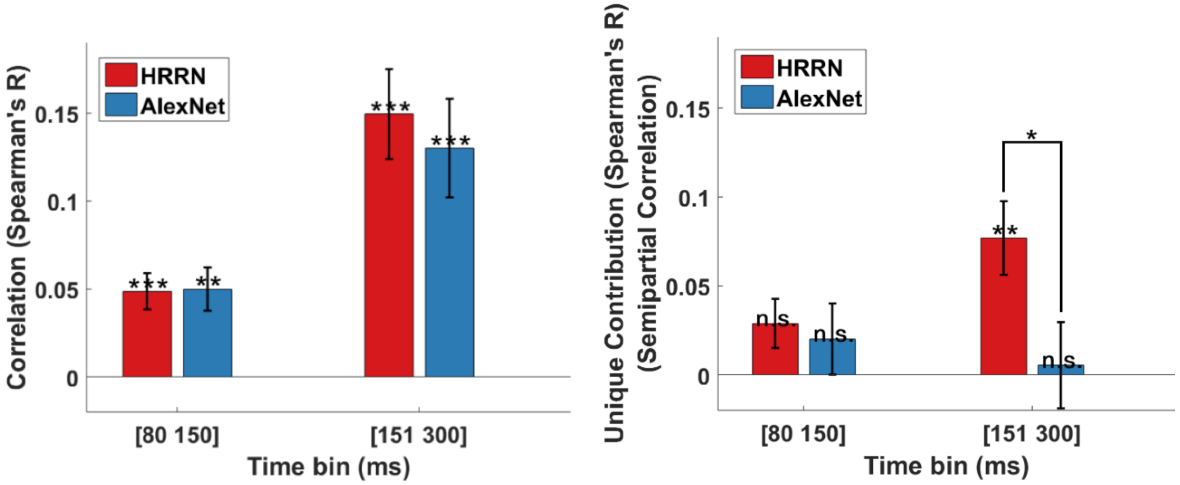


**Figure S9. Contribution of the feedforward and recurrent models in explaining MEG data under 0% occlusion.** (a) Correlation between the models RDMs and the average MEG RDM over two different time bins. (b) Unique contribution of each model (semipartial correlation) in explaining the MEG data. Error bars represent SEM (Standard Error of the Mean). Significantly above zero correlations/semipartial-correlations and significant differences between the two models are indicated by stars. * = $p<0.05$ ; ** = $p<0.01$; *** = $p<0.001$.

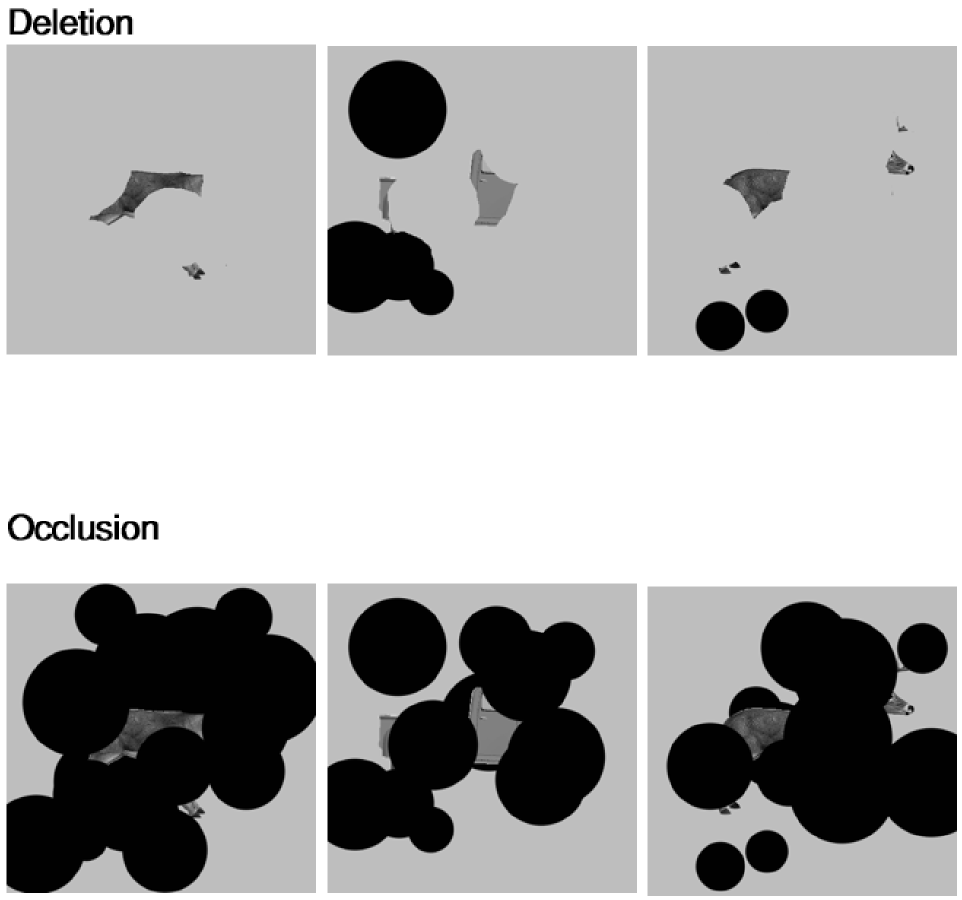


**Figure S10. Sample images of object occlusion versus deletion.**

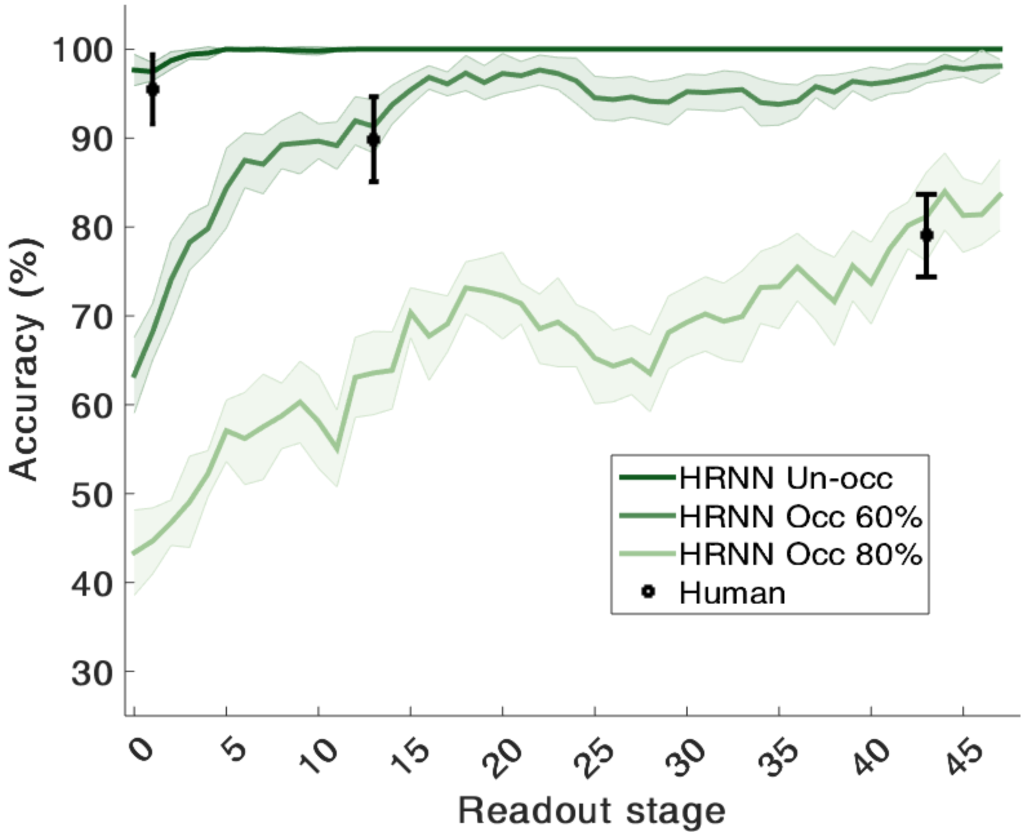


**Figure S11. HRRN accuracy across readout stages for different levels of occlusion.** Shaded error bars indicate SD. Black circles are average accuracies across n = 16 human participants. Readout stage: readout stage refers to the number of local recurrent iterations involved in processing the input image throughout the hierarchy of the network. Readout stage 0 is when the model is fully feedforward (no local recurrent is active). And readout stage 1 is when only one recurrent iteration is engaged and readout stage *n* is when the network has gone through *n* recurrent iterations.

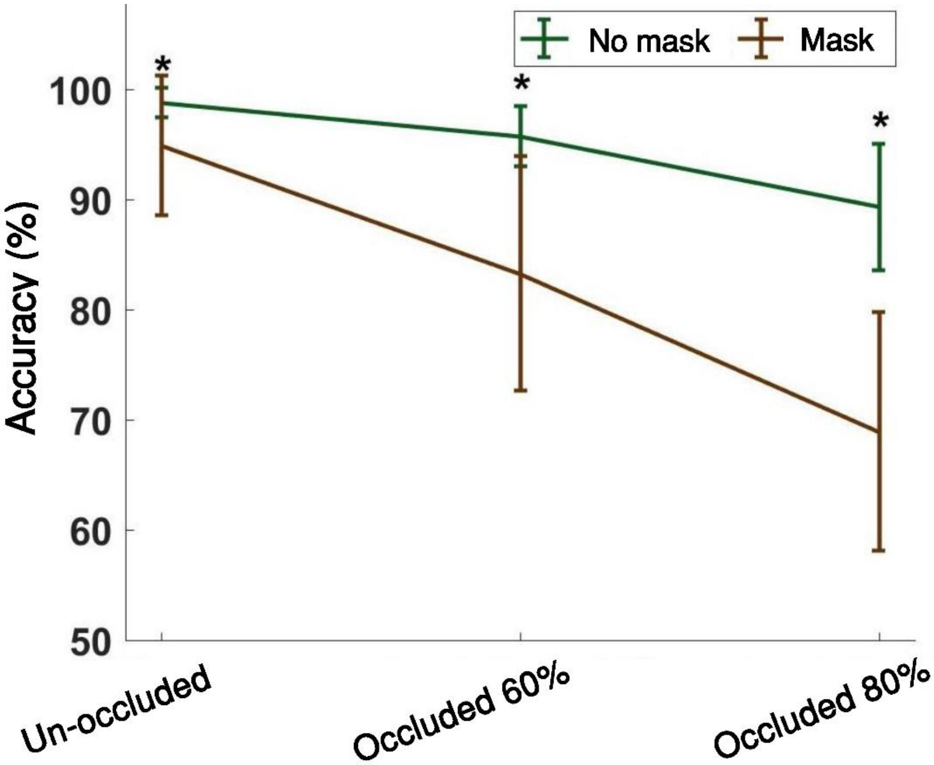


**Figure S12. Behavioral performance of animate/inanimate categorization task of the MEG experiment.** Stars indicate significant differences between mask and no-mask trials. The results are averaged over N=15 human participants.

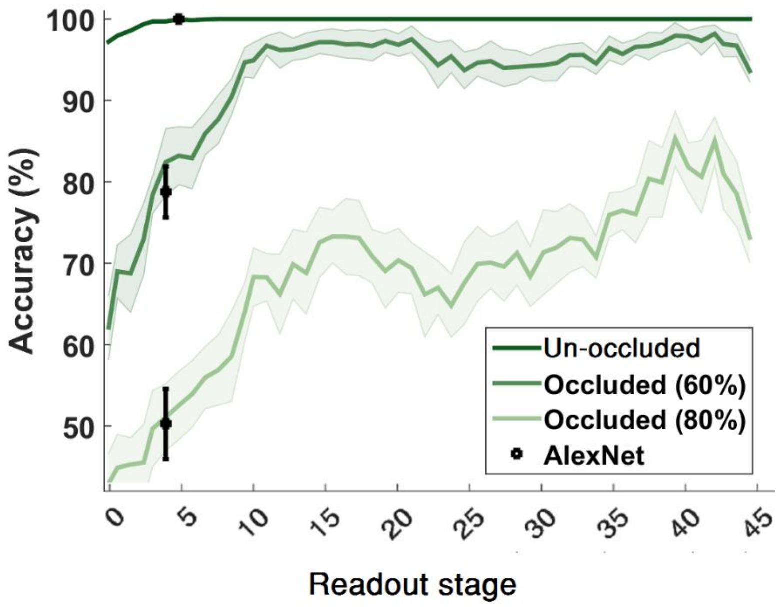


**Figure S13. Categorization accuracies of HRRN across different readout stages compared with Alexnet for different occlusion levels.** Shaded error bars indicate SD. Readout stage: readout stage refers to the number of local recurrent iterations involved in processing the input image throughout the hierarchy of the network. Readout stage 0 is when the model is fully feedforward (no local recurrent is active). And readout stage 1 is when only one recurrent iteration is engaged and readout stage n is when the network has gone through n recurrent iterations. Black circles are average accuracies for Alexnet, which are shown around the approximate corresponding HRRN readout stages.

**Sensorwise visualization of pairwise object decoding across time for no-occlusion (Movie 1) and occlusion (Movie 2) conditions.** Color map represents percent of decoding accuracies across head surface (chance level = 50%). Circles indicate neighboring triplets (102 triplets) of MEG sensors (2 gradiometers and 1 magnetometer in each location). Significantly above chance decoding accuracies after correction for multiple comparison are shown by black dots (FDR-corrected across 102 triplets and 801 time-points). Gray dots indicate decoding accuracies with p<0.05 (right sided signed rank test) that did not remain signifiant after FDR-correction. At each time point, the peak decoding accuracy is indicated by a red dot.
